# Low-level repressive histone marks fine-tune stemness gene transcription in neural stem cells

**DOI:** 10.1101/2022.11.18.517130

**Authors:** Arjun Rajan, Lucas Anhezini, Noemi Rives-Quinto, Megan C. Neville, Elizabeth D. Larson, Stephen F. Goodwin, Melissa M. Harrison, Cheng-Yu Lee

## Abstract

Coordinated regulation of stemness gene activity by transcriptional and translational mechanisms poise stem cells for a timely cell-state transition during differentiation. Although important for all stemness-to-differentiation transitions, mechanistic understanding of the fine-tuning of stemness gene transcription is lacking due to the compensatory effect of translational control. We used intermediate neural progenitor (INP) identity commitment to define the mechanisms that fine-tune stemness gene transcription in fly neural stem cells (neuroblasts). We demonstrate that the transcription factor Fruitless^C^ (Fru^C^) binds *cis*-regulatory elements of most genes uniquely transcribed in neuroblasts. Loss of *fru^C^* function alone has no effect on INP commitment but drives INP dedifferentiation when translational control is reduced. Fru^C^ negatively regulates gene expression by promoting low-level enrichment of the repressive histone mark H3K27me3 in gene *cis*-regulatory regions. Identical to *fru^C^* loss-of-function, reducing Polycomb Repressive Complex 2 activity increases stemness gene activity. We propose low-level H3K27me3 enrichment fine-tunes stemness gene transcription in stem cells, a mechanism likely conserved from flies to humans.

## Introduction

Expression of genes that promote stemness or differentiation must be properly controlled in stem cells to allow their progeny to transition through various intermediate stages of cell fate specification in a timely fashion (Bhaduri et al., 2021; Dillon et al., 2022; Michki et al., 2021; Pollen et al., 2015; Ruan et al., 2021). Exceedingly high levels of stemness gene transcripts that promote an undifferentiated state in stem cells can overwhelm translational control that downregulates their activity in stem cell progeny and perturb timely onset of differentiation (Larson et al., 2021; Ohtsuka and Kageyama, 2021; San-Juán and Baonza, 2011; Xiao et al., 2012; Zacharioudaki et al., 2012; Zhu et al., 2012). Conversely, excessive transcription of differentiation genes that instill biases toward terminal cellular functions in stem cells can overcome the mechanisms that uncouple these transcripts from the translational machinery and prematurely deplete the stem cell pool (Baser et al., 2019; de Rooij et al., 2019; Lennox et al., 2018). Thus, fine-tuning stemness and differentiation gene transcription in stem cells minimizes inappropriate gene activity that could result in developmental anomalies. Coordinated regulation of stemness and differentiation gene activity in stem cells by transcriptional and translational control poise stem cell progeny for a timely cell-state transition during differentiation (Ables et al., 2011; Bigas and Porcheri, 2018; Kobayashi and Kageyama, 2014; Koch et al., 2013; Rajan et al., 2021). Mechanistic investigation of the fine-tuning of stemness and differentiation gene transcription *in vivo* is challenging due to the compensatory effect of translational control, a lack of sensitized functional readouts, and a lack of insight into relevant transcription factors.

Neuroblast lineages of the fly larval brain provide an excellent *in vivo* paradigm for mechanistic investigation of gene regulation during developmental transitions because the cell-type hierarchy is well-characterized at functional and molecular levels (Doe, 2017; Homem et al., 2015; Janssens and Lee, 2014). A larval brain lobe contains approximately 100 neuroblasts, and each neuroblast asymmetrically divides every 60-90 minutes, regenerating itself and producing a sibling progeny that commits to generating differentiated cell types. Most of these neuroblasts are type I, which generate a ganglion mother cell (GMC) in every division. A GMC undergoes terminal division to produce two neurons. Eight neuroblasts are type II, which invariably generate an immature intermediate neural progenitor (immature INP) in every division (Bello et al., 2008; Boone and Doe, 2008; Bowman et al., 2008). An immature INP initiates INP commitment 60 minutes after asymmetric neuroblast division (Janssens et al., 2017). An immature INP initially lacks Asense (Ase) protein expression and upregulates Ase as it progresses through INP commitment. Once INP commitment is complete, an Ase^+^ immature INP transitions into an INP and asymmetrically divides 5-6 times to generate more than a dozen differentiated cells, including neurons and glia (Bayraktar and Doe, 2013; Viktorin et al., 2011). All type II neuroblast lineage cell types can be unambiguously identified based on functional characteristics and protein marker expression in larval brains. Single-cell RNA-sequencing (scRNA-seq) of sorted, fluorescently labeled INPs and their differentiating progeny from wild-type brain tissue has led to the discovery of many new genes that contribute to the generation of diverse differentiated cell types during neurogenesis (Michki *et al*., 2021). This wealth of information on the type II neuroblast lineage allows for mechanistic investigations of precise spatiotemporal regulation of gene expression during developmental transitions.

A multilayered gene regulation system ensures timely onset of INP commitment in immature INPs by coordinately terminating Notch signaling activity (Komori et al., 2018). Activated Notch signaling drives the expression of downstream-effector genes *deadpan (dpn)* and *Enhancer of (splitz)mγ* (*E(spl)mγ*), which promote stemness in type II neuroblasts by poising activation of the master regulator of INP commitment *earmuff* (*erm*) (San-Juán and Baonza, 2011; Xiao *et al*., 2012; Zacharioudaki et al., 2016; Zacharioudaki *et al*., 2012; Zhu *et al*., 2012). During asymmetric neuroblast division, the basal protein Numb and Brain tumor (Brat) exclusively segregate into immature INPs, where they terminate Notch signaling activity and promote the timely onset of Erm expression (Bello et al., 2006; Betschinger et al., 2006; Lee et al., 2006a; Lee et al., 2006c; Wang et al., 2006). Numb is a conserved Notch inhibitor and prevents continual Notch activation in immature INPs (Frise et al., 1996; Lee *et al*., 2006a; Wang *et al*., 2006; Wirtz-Peitz et al., 2008; Zhong et al., 1996). Asymmetric segregation of the RNA-binding protein Brat is facilitated by its adapter protein Miranda, which releases Brat from the cortex of immature INPs, allowing Brat to promote decay of Notch downstream-effector gene transcripts and thus initiate differentiation (Komori *et al*., 2018; Laver et al., 2015; Loedige et al., 2015; Loedige et al., 2014; Reichardt et al., 2018). Complete loss of *numb* or *brat* function leads to unrestrained activation of Notch signaling in immature INPs driving them to revert into type II neuroblasts leading to a severe supernumerary neuroblast phenotype. Similarly, increased levels of activated Notch or Notch transcriptional target gene expression in immature INPs can drastically enhance the moderate supernumerary neuroblast phenotype in *brat*- or *numb*-hypomorphic brains (Janssens et al., 2014; Komori *et al*., 2018; Komori et al., 2014b; Larson *et al*., 2021; Xiao *et al*., 2012). Collectively, these findings suggest that precise transcriptional control of *Notch* and Notch target gene expression levels during asymmetric neuroblast division is essential, safeguarding the generation of neurons that are required for neuronal circuit formation in adult brains.

We defined the fine-tuning of stemness gene transcription as a function that is mild enough to not effect INP commitment when lost alone but enough to enhance immature INP reversion to supernumerary neuroblasts induced by decreased post-transcriptional control of stemness gene expression. We established three key criteria to identify regulators which fine-tune stemness gene transcription in neuroblasts, (1) an established role in transcriptional regulation, for example a DNA-binding transcription factor and (2) clear expression in neuroblasts with no protein expression in immature INPs, (3) acts as a negative regulator of its targets. From a type II neuroblast lineage-specific single-cell gene transcriptomic atlas, we found that *fruitless* (*fru*) mRNAs are detected in type I & II neuroblasts but not in their differentiating progeny. One specific Zn-finger containing isoform of Fru (Fru^C^) is exclusively expressed in all neuroblasts. Fru^C^ binds *cis*-regulatory elements of most genes uniquely transcribed in type II neuroblasts, including *Notch* and Notch downstream-effector genes that promote stemness in neuroblasts. A modest increase in *Notch* or Notch downstream gene expression induced by loss of *fru^C^* function alone has no effect on INP commitment, but enhances immature INP reversion to type II neuroblasts in *numb*- and *brat*-hypomorphic brains. To establish how Fru^C^ might fine-tune gene transcription in neuroblasts, we examined the distribution of established histone modifications in the presence or absence of *fru^C^*. We surprisingly found Fru^C^-dependent low-level enrichment of the repressive histone marker H3K27me3 in most Fru^C^-bound peaks in genes uniquely transcribed in type II neuroblasts including *Notch* and its downstream-effector genes. The Polycomb Repressive Complex 2 (PRC2) subunits are enriched in FruC-bound peaks in genes uniquely transcribed in type II neuroblasts, and reduced PRC2 function enhances the supernumerary neuroblast phenotype in *numb*-hypomorphic brains, identical to *fru^C^* loss-of-function. We conclude that the Fru^C^-PRC2-H3K27me3 molecular pathway fine-tunes stemness gene expression in neuroblasts by promoting low-level H3K27me3 enrichment in their *cis*-regulatory elements. The mechanism by which PRC2-H3K27me3 fine-tune stem cell gene expression will likely be relevant throughout metazoans.

## Results

### A gene expression atlas captures dynamic changes throughout type II neuroblast lineages

To identify regulators of gene transcription during asymmetric neuroblast division, we constructed a single-cell gene transcription atlas that encompasses all cell types in the type II neuroblast lineage in larval brains. We fluorescently labeled all cell types in the lineage in wild-type third-instar larval brains, sorted positively labeled cells by flow cytometry, and performed single-cell RNA-sequencing (scRNA-seq) using a 10X genomic platform (Fig. 1A, S1A). This new dataset displays high levels of correlation to our previously published scRNA-seq dataset which were limited to INPs and their progeny. The harmonization of these two datasets results in a gene transcription atlas of the type II neuroblast lineage consisting of over 11,000 cells (Fig. 1B). Based on the expression of known cell identity genes, we were able to predict clusters consisting of type II neuroblasts (*dpn^+^,pnt^+^*), INPs (*dpn^+^,opa^+^*), GMCs (*dap^+^,hey^-^)*, immature neurons (*dap^+^,hey^+^*), mature neurons (*hey^-^,nSyb^+^*), and glia (*repo^+^*) (Fig. 1C, S1B). The UMAP positions of these clusters match well with the results of pseudo-time analyses from a starter cell that was positive for *dpn*, *pnt*, and *RFP* transcripts (Fig. 1D). Thus, the harmonized scRNA-seq dataset captures all molecularly and functionally defined stages of differentiation in the type II neuroblast lineage (Fig. 1E).

**Fig. 1.**
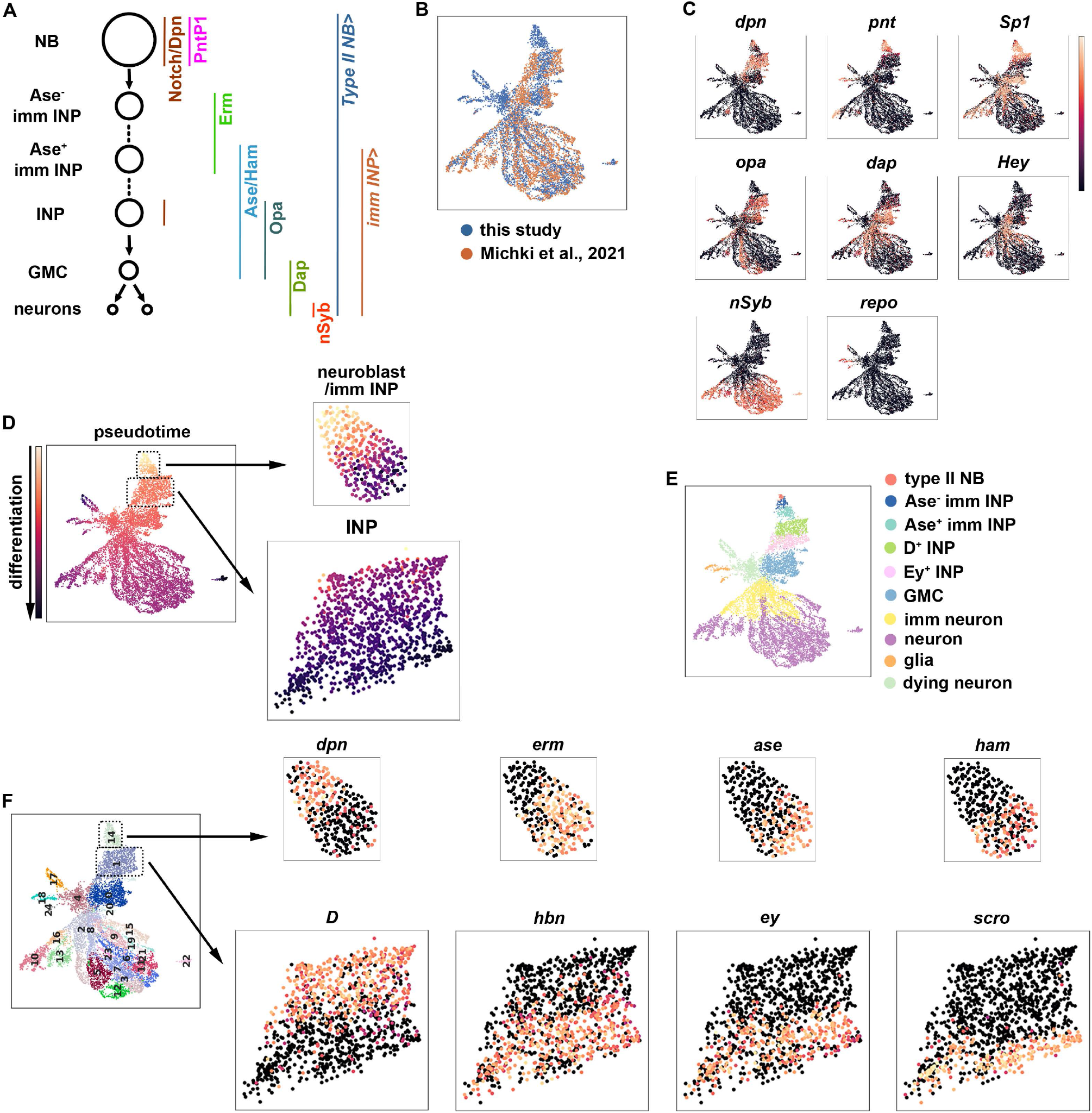
A single-cell gene expression atlas of type II neuroblast lineages. (A) Summary illustration of the type II neuroblast lineage and the Gal4 drivers used for sorting cell types in this lineage from wild-type larval brains for scRNA-seq. (B) Harmonization of the scRNA-seq dataset from the entire type II neuroblast (NB) lineage generated in this study (blue) and our previously published scRNA-seq dataset which were limited to INPs and their progeny (orange). (C) UMAPs of known celltype-specific marker genes. (D) Pseudotime analysis starting from genes enriched for *dpn*, *pnt*, and *RFP* transcripts. (E) Annotated gene expression atlas of a wild-type type II neuroblast lineage. (F) Left: Leiden clustering of the scRNA-seq atlas. Right: Representative UMAPs of dynamic transcription factors for clusters 14 (NBs and immature INPs) and 1 (INPs).

To determine whether the new scRNA-seq dataset encompasses neuroblast progeny undergoing dynamic changes in cell identity during differentiation, we examined transcripts that were transiently expressed in neuroblast progeny undergoing INP commitment or asymmetric INP division. We found that cluster 14 contains type II neuroblasts (*dpn^+^,erm^-^,ase^-^,ham^-^*), Ase^-^ immature INPs (*dpn^-^,erm^+^,ase^-^,ham^-^*) and Ase^+^ immature INPs (*dpn^-^,erm^-^,ase^+^,ham^+^*) (Fig. 1F). Furthermore, cluster 1 contains proliferating INPs including young INPs (*D^+^,hbn^-^,ey^-^,scro^-^*) and old INPs (*D^-^,hbn^-^,ey^+^,scro^+^*) (Fig. 1F). These data led us to conclude that the type II neuroblast lineage gene transcription atlas captures neuroblast progeny undergoing dynamic changes in cell identity during differentiation (Fig. 1E).

### FruC negatively regulates stemness gene expression in neuroblasts

We hypothesized that regulators that fine-tune stemness gene expression in neuroblasts should (1) be transcription factors, (2) be exclusively expressed in type II neuroblasts, and (3) negatively regulate gene transcription. We searched for candidate genes that fulfill these criteria in the cluster 14 of the type II neuroblast lineage gene transcription atlas. *dpn* serves as a positive control because its transcripts are highly enriched in type II neuroblasts and rapidly degraded in Ase^-^ immature INPs, allowing us to distinguish neuroblasts from immature INPs (Fig. 2A–B). We found the expression of *fru* mirrors *dpn* expression, with transcript levels high in type II neuroblasts but lower in Ase^-^ immature INPs (Fig. 2A). *fru* is a pleiotropic gene with at least two major functions: one that controls male sexual behavior and another that is essential for viability in both sexes (Goodwin and Hobert, 2021). *fru* transcripts are alternatively spliced into multiple isoforms that encode putative transcription factors containing a common BTB (protein-protein interaction) N-terminal domain and one of four C-terminal zinc-finger DNA-binding domains (Dalton et al., 2013; Neville et al., 2014; von Philipsborn et al., 2014) (Fig. 2A). We used the Fru-common antibody that recognizes all isoforms to determine the spatial expression pattern of Fru protein in green fluorescent protein (GFP)-marked wild-type neuroblast clones. We detected Fru in neuroblasts but found that Fru is rapidly downregulated in their differentiating progeny in type I and II lineages (Fig. 2B). To determine which Fru isoform is expressed in neuroblasts, we examined the expression of isoform-specific *fru::Myc* tagged alleles (von Philipsborn *et al*., 2014). We found that Fru^C^::Myc is specifically expressed in both types of neuroblasts but not in their differentiating progeny, including immature INPs, INPs, and GMCs (Fig. 2C). These data indicate that Fru^C^ is the predominant Fru isoform expressed in neuroblasts.

**Fig. 2.**
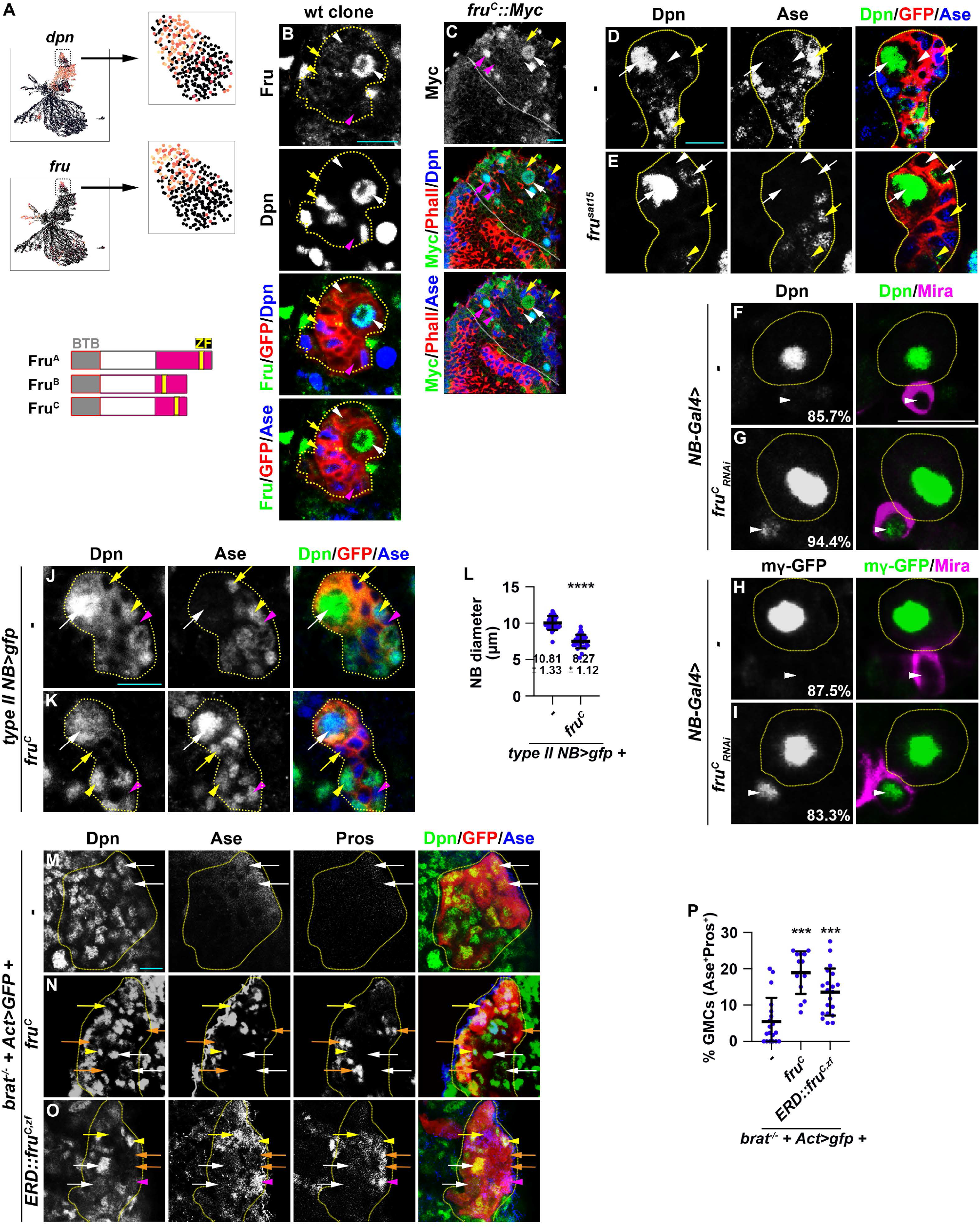
Fru^c^ functions through transcriptional repression to regulate stemness gene expression. (A) <u>Top</u>: *fru* and *dpn* mRNAs are detected in neuroblasts in cluster 14 of the scRNA-seq dataset, but only *dpn* mRNAs are detected in INPs in cluster 1. <u>Bottom</u>: Domains in Fru protein isoforms. (B) Fru protein is detected in the neuroblast but not in INPs in a GFP-marked type II neuroblast lineage clone. (C) Endogenously expressed Fru^c^::Myc is detected in type I & II neuroblasts but not in their differentiating progeny. White dotted line separates optic lobe from brain. (D-E) Dpn is ectopically expressed in Ase^-^ immature INPs in a GFP-marked mosaic clone derived from a single *fru*-null (*fru^sat15^*) type II neuroblast. (F-I) Reducing *fru^c^* function leads to a high frequency of ectopic Dpn and E(spl)mγ::GFP expression in the newborn immature INP marked by cortical Miranda staining. (J-L) Type II neuroblasts overexpressing Fru^c^ display differentiation markers including a reduced cell diameter and Ase expression. Quantification of the cell diameter is shown in L. (M-P) Overexpressing full-length Fru^c^ or a constitutive transcriptional repressor form of Fru^c^ (Fru^c,zf^::ERD) restores differentiation in *brat*-null type II neuroblast clones. The percentage of GMCs per clone is shown in P. The following labeling applies to all images in this paper: yellow dashed line encircles a type II neuroblast lineage; white arrow: type II neuroblast; white arrowhead: Ase^-^ immature INP; yellow arrow: Ase^+^ immature INP; yellow arrowhead: INP; magenta arrow: type I neuroblast; magenta arrowhead: GMC; orange arrowhead: neurons. Scale bars: 10 μm. P-value: ***<0.0005, and ****<0.00005

To define the function of Fru in neuroblasts, we assessed the identity of cells in the GFP-marked mosaic clones derived from single type II neuroblasts. The wild-type neuroblast clone always contains a single neuroblast that can be uniquely identified by cell size (10-12 μm in diameter) and marker expression (Dpn^+^Ase^-^) as well as 6-8 smaller, Dpn^-^ immature INPs (Fig. 2D). The neuroblast clone carrying deletion of the *fru* locus (*fru^-/-^*) contains a single identifiable neuroblast but frequently contains multiple Ase^-^ immature INPs with detectable Dpn expression (Fig. 2E). Over 80% of newborn immature INPs (marked by intense cortical Mira expression) generated by *fru^C^*-mutant type II neuroblasts ectopically express Dpn and E(spl)m*γ* while greater than 85% of newborn immature INPs generated by wild-type neuroblasts do not express these Notch downstream-effectors (Fig. 2F–I). These results support a model in which loss of *fru^C^* function increases the expression of Notch downstream-effectors genes that promote stemness in neuroblasts. Type II neuroblasts overexpressing Fru^C^ prematurely initiate INP commitment, as indicated by a reduced cell diameter and precocious Ase expression (Fig. 2J–L). Thus, loss of *fru^C^* function increases stemness gene expression whereas gain of *fru^C^* decreases stemness gene expression during asymmetric neuroblast division.

To determine whether Fru^C^ negatively regulates the transcription of stemness genes in type II neuroblasts, we overexpressed wild-type Fru^C^ in GFP-marked neuroblast lineage clones in *brat*-null brains. Ectopic translation of Notch downstream-effector gene transcripts that promote stemness in neuroblasts drives immature INP reversion to supernumerary type II neuroblasts at the expense of differentiating cell types in *brat*-null brains (Komori *et al*., 2018; Loedige *et al*., 2015; Reichardt *et al*., 2018). Control clones in *brat*-null brains contain mostly type II neuroblasts and few differentiating cells that include Ase^+^ immature INPs, INPs, and GMCs (Fig. 2M, 2P). By contrast, overexpressing full-length Fru^C^ increases the number of INPs, GMCs, and differentiating neurons (Ase^-^Pros^+^) in *brat*-null neuroblast clones (Fig. 2N, 2P). This result indicates that Fru^C^ overexpression is sufficient to restore differentiation in *brat*-null brains. Similar to full-length Fru^C^ overexpression, overexpressing a constitutive repressor form of Fru^C^ consisting of the <u>E</u>ngrail <u>R</u>epressor <u>D</u>omain fused in frame with the zinc-finger DNA-binding motif of Fru^C^ (Fru^C,zf^::ERD) also restored differentiation in *brat*-null brains (Fig. 2O–P). Thus, we conclude that Fru^C^ negatively regulates stemness gene expression in type II neuroblasts.

### Fru^c^ binds *cis*-regulatory elements of Notch pathway genes essential for neuroblast stemness

If Fru^C^ directly represses stemness gene expression, Fru^C^ should bind their *cis*-regulatory elements. To identify Fru^C^-bound regions in neuroblasts, we applied a protocol of Cleavage Under Targets and Release Using Nuclease (CUT&RUN) to brain extracts from dissected third-instar *brat*-null larvae homozygous for the *fru^C^::Myc* allele. *brat*-null brains accumulate thousands of supernumerary type II neuroblasts at the expense of INPs and provide a biologically relevant source of type II neuroblast-specific chromatin (Janssens *et al*., 2017; Komori *et al*., 2018; Komori et al., 2014a; Larson *et al*., 2021; Rives-Quinto et al., 2020). We used a specific antibody against the Myc epitope or the Fru-common antibody to confirm that Fru^C^::Myc is detected in all supernumerary type II neuroblasts in *brat*-null brains homozygous for *fru^C^::Myc* (Fig. S2A). We determined the genome-wide occupancy of Fru^C^::Myc in type II neuroblasts using the Myc antibody and Fru-common antibody, and found that Fru^C^::Myc binding patterns revealed by these two antibodies are highly correlated (Fig. S2B; Pearson correlation = 0.94). Fru^C^ binds 9301 regions in type II neuroblasts (Fig. 3A). Overall, 59% of Fru^C^-bound regions are promoters whereas 29% are enhancers in the intergenic and intronic regions (Fig. 3A). By contrast, 15% of randomized control regions are promoters and 55% are enhancers (Fig. 3A). 50.1% of Fru^C^-bound regions in promoters and enhancers overlap with regions of accessible chromatin (Larson *et al*., 2021) (Fig. 3B). Consistent with the finding that Fru^C^ negatively regulates stemness gene expression, Fru^C^ binds promoters and neuroblast-specific enhancers of *Notch, dpn, E(spl)mγ, klumpfuss* (*klu*) and *tailless* (*tll*) that were previously shown to maintain type II neuroblasts in an undifferentiated state (Fig. 3C, Fig. S2C). 74% of genes uniquely transcribed in type II neuroblasts (NB genes) are bound by Fru^C^ whereas 41% of these genes are in randomized control (Fig. 3D–E). By contrast, the percentage of Fru^C^-bound genes transcribed throughout the type II neuroblast lineage is similar to random control (Fig. 3D–E). Because stemness gene transcripts are highly enriched in type II neuroblasts, these results suggest that Fru^C^ preferentially binds *cis*-regulatory elements of stemness genes.

**Fig. 3.**
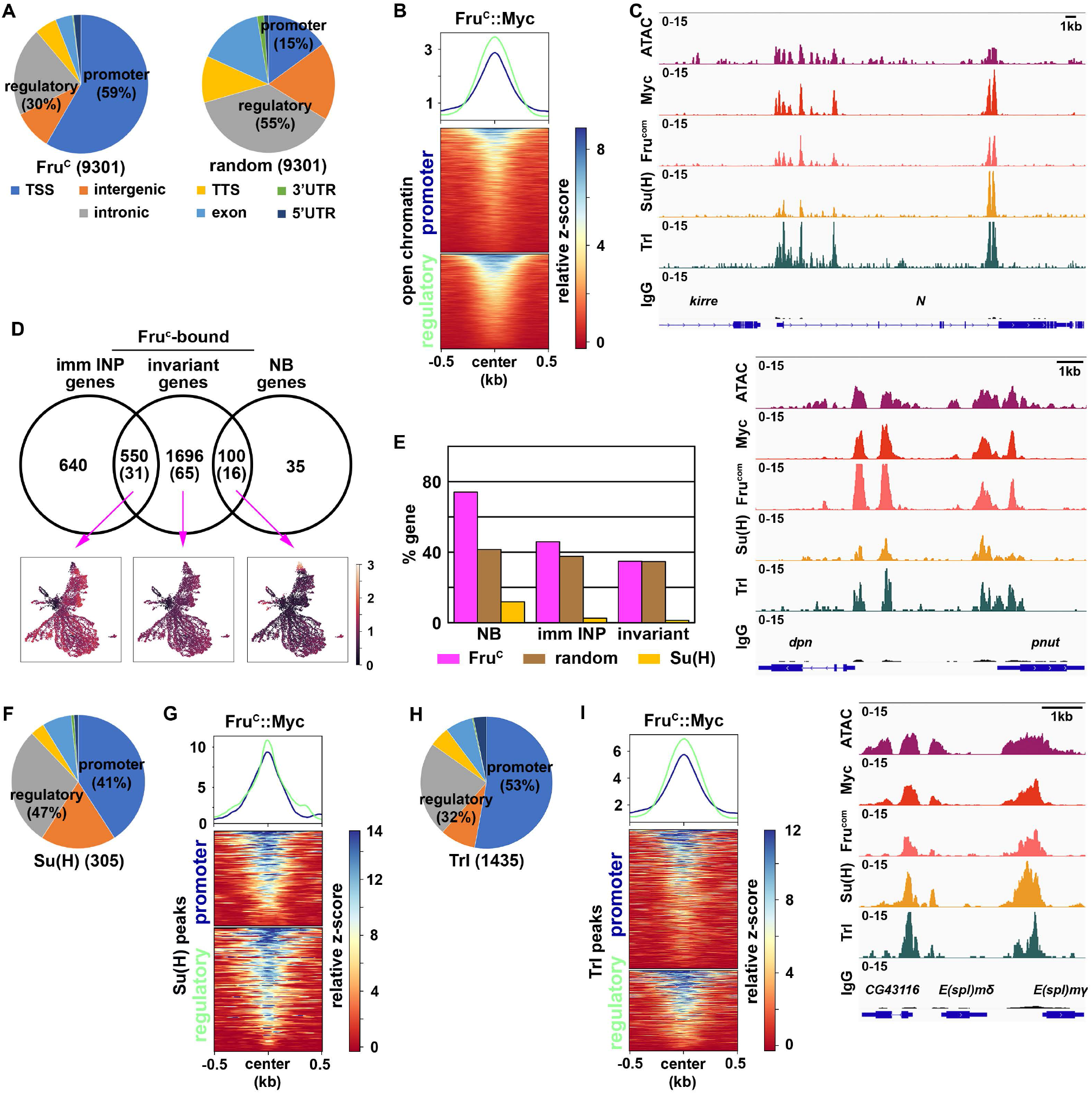
Fru^c^ preferentially binds regulatory elements of genes uniquely expressed in neuroblasts. (A) Genomic binding distribution of Fru^C^-bound peaks (total # of peaks shown in parentheses) from CUT&RUN or random (Fru^c^ shuffled) in type II neuroblast-enriched chromatin from *brat-*null brains show that Fru^c^ preferentially binds promoters. (B) Heatmap is centered on promoters or regulatory regions showing accessible chromatin as defined by ATAC-seq with 500-bp flanking regions and ordered by signal intensity of Fru^c^::Myc binding. (C) Representative z score-normalized genome browser tracks showing regions with accessible chromatin (ATAC-seq) and bound by Fru^c^ (detected by the Myc antibody or Fru^COM^ antibody), Su(H), Trl, or IgG at *Notch, dpn*, and *E(spl)mγ* loci. (D-E) Genes in cluster 14 from the scRNA-seq dataset were separated into neuroblast-enriched genes (right), immature INP-enriched genes (left), and invariant genes. The middle circle is the set of genes bound by Fru^c^ or Su(H) (shown in parentheses). UMAPs show gene enrichment score for Fru^c^-bound genes uniquely expressed in neuroblasts (right), uniquely expressed in immature INPs (left), and ubiquitously expressed (invariant genes) (middle). (F-G) Genomic binding distribution of identified Su(H)-bound peaks (total # of peaks shown in parentheses) from CUT&RUN in type II neuroblast-enriched chromatin. Heatmap is centered on promoters or regulatory regions and ordered by signal intensity of Su(H) binding. (H-I) Genomic binding distribution of identified Trl-bound peaks (total # of peaks shown in parentheses) from CUT&RUN in type II neuroblast-enriched chromatin. Heatmap is centered on promoters or regulatory regions and ordered by signal intensity of Trl binding.

A mechanism by which Fru^C^ can negatively regulate stemness gene expression levels is to modulate the activity of the Notch transcriptional activator complex activity. Using Affymetrix GeneChip, a previous study demonstrated that the Notch transcriptional activator complex binds 595 regions in 185 transcribed genes in neuroblasts (log_2_ FC>0.5) including *dpn*, *E(spl)mγ, klu*, and *tll* (Zacharioudaki *et al*., 2016). To precisely identify Notch-bound peaks in type II neuroblasts, we used a specific antibody against the DNA-binding subunit of the Notch transcriptional activator complex, Suppressor of Hairless (Su(H)), to perform a CUT&RUN assay on brain lysate form third-instar *brat*-null larvae homozygous for *Fru^C^::Myc*. Prior to the genomic study, we validated the specificity of the Su(H) antibody *in vivo* by performing immunofluorescent staining of *brat*-null larvae homozygous for *fru^C^::Myc*. We observed co-expression of Su(H) and Dpn in thousands of supernumerary type II neuroblasts in *brat*-null brains (Fig. S2D). Because *dpn* is directly activated by Notch in many cell types, including neuroblasts, this result confirms that the Su(H) antibody can detect activated Notch activity. We found that Su(H) binds 305 regions in 112 genes in type II neuroblasts, and that Su(H)-bound regions are predominantly in promoters and enhancers (Fig. 3F). Overall, 95.2% of Su(H)-bound promoters and 90.8% of Su(H)-bound enhancers overlap with Fru^C^-bound peaks (Fig. 3G). These peaks include the promoters of *Notch, dpn, E(spl)mγ, klu* and *tll* as well as the enhancers that drive their expression in neuroblasts (Fig. 3C, Fig. S2C). Our data support a model that Fru^C^ regulates Notch pathway gene expression by occupying functionally relevant regulatory elements bound by Notch in type II neuroblasts.

*De novo* motif discovery identified a sequence bound by the transcription factor Trithorax-like (Trl), also known as GAGA factor, is significantly enriched in both Fru^C^-bound promoters and enhancers (Fig. S2E). The Trl motif was previously found to be the most significantly enriched in Fru^C^-associated genomic regions in the larval nervous system (Neville *et al*., 2014). Trl, like Fru, is a member of the BTB-Zn-finger transcription factor family which heterodimerize with other BTB-domain-containing factors to regulate gene transcription (Bonchuk et al., 2022). Trl is an evolutionarily conserved multifaceted transcription factor that regulates diverse biological processes by interacting with a wide variety of proteins including PRC2 complex components (Chetverina et al., 2021; Lomaev et al., 2017; Srivastava et al., 2021).

Although the Trl motif is generally enriched at promoters, enrichment of this motif in Fru^C^-bound promoters and enhancers suggest that Fru^C^ might function together with Trl to negatively regulate gene transcription in type II neuroblasts. We examined whether Trl indeed binds Fru^C^-bound regions in type II neuroblasts using a specific antibody against Trl (Judd et al., 2021). We validated the specificity of the Trl antibody *in vivo* by performing immunofluorescent staining of *brat*-null larvae homozygous for *fru^C^::Myc*. We found that Trl is highly enriched in the nuclei of thousands of supernumerary type II neuroblasts marked by Dpn expression (Fig. S2F). We used the Trl antibody to perform a CUT&RUN assay on brain lysate from third-instar *brat*-null larvae homozygous for *fru^C^::Myc*. We identified 1435 Trl-bound regions, including promoters and enhancers, in type II neuroblasts (Fig. 3H). In total, 85.4% of Trl-bound promoters and 87.8% of Trl-bound enhancers overlapped with Fru^C^-bound regions including *Notch, dpn, E(spl)mγ, klu* and *tll* loci (Fig. 3C, 3I, S2C). These data suggest that Fru^C^ may function together with Trl to regulate the transcription of stemness genes in neuroblasts.

### Fru^C^ fine-tunes the expression of Notch pathway genes during asymmetric neuroblast division

Loss- and gain-of-function of *fru^C^* mildly alters the expression of Notch downstream-effector genes that promote stemness in neuroblasts (Fig. 2). Fru^C^ occupies enhancers relevant to the functions of Notch downstream-effector genes in type II neuroblasts (Fig. 3). Thus, Fru^C^ is an excellent candidate for finetuning stemness gene expression in neuroblasts. To functionally validate the role of Fru^C^ in fine-tuning stemness gene expression, we tested whether loss of *fru^C^* function can enhance the supernumerary neuroblast phenotype in *brat*-hypomorphic (*brat^hypo^*) brains. Immature INPs revert to supernumerary type II neuroblasts at low frequency due to a modest increase in Notch downstream-effector gene expression in *brat^hypo^* brains (Komori *et al*., 2018). Consistent with the finding that reduced *fru* function increases Notch downstream-effector protein levels in immature INPs, the heterozygosity of a *fru* deletion (*fru^-/+^*) enhances the supernumerary neuroblast phenotype in *brat^hypo^* brains (Fig. 4A). Furthermore, *brat^hypo^* brains lacking *fru^C^* function displayed greater than a two-fold increases in supernumerary neuroblasts compared with *brat^hypo^* brains heterozygous for a *fru* deletion (Fig. 4A). These data support our model that Fru^C^ fine-tunes the expression of Notch downstream-effector genes that promote stemness in neuroblasts.

**Fig. 4.**
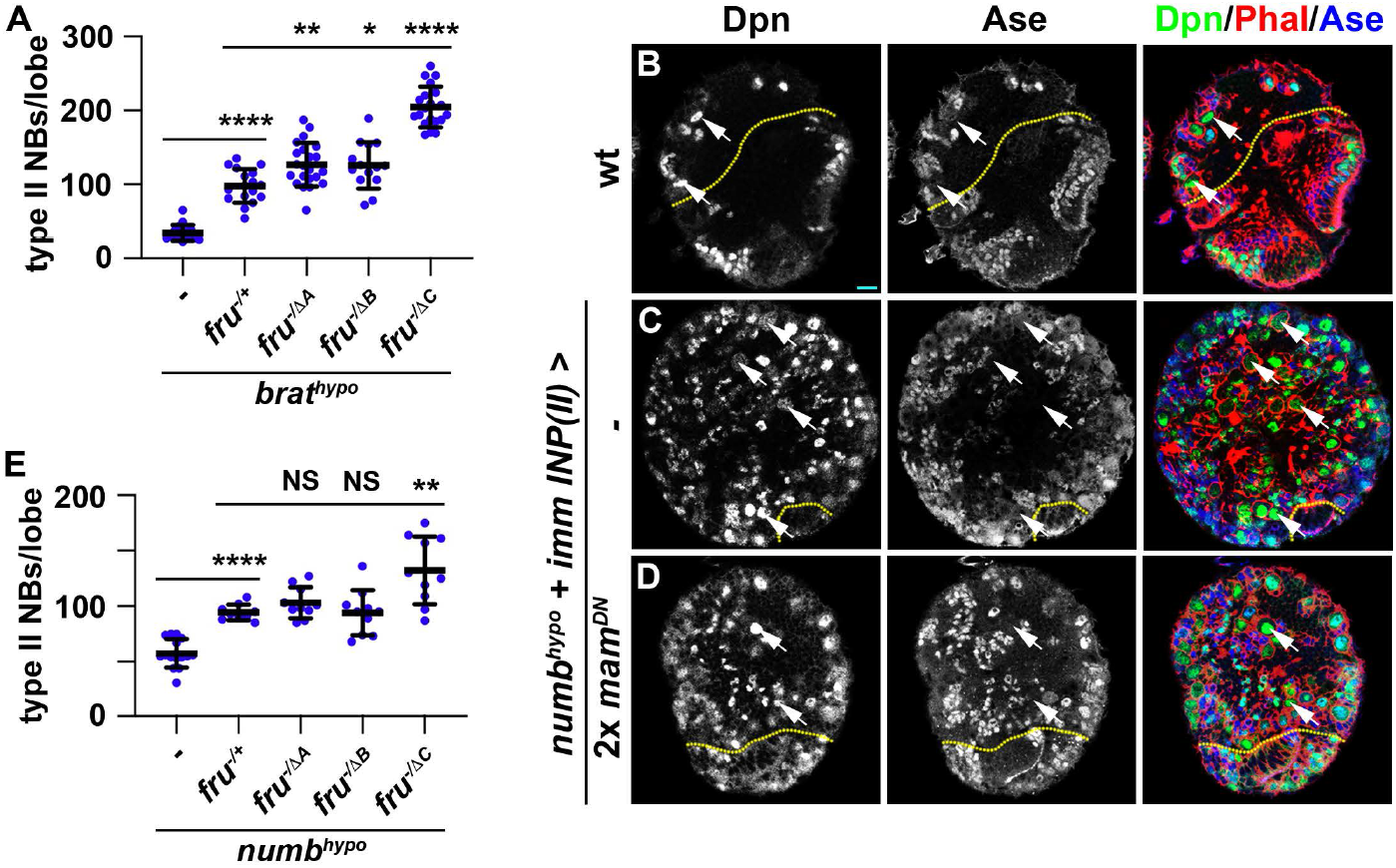
Loss of *fru^C^* function enhances stemness gene activity in *brat^hypo^* and *numb^hypo^* brains. (A) Loss of *fru^C^* function (*fru^△C/-^*) enhances the supernumerary neuroblast phenotype in *brat^hypo^* brains heterozygous for a *fru* deletion (*fru^-/+^*), but loss of *fru^A^* (*fru^△A/-^*) or *fru^B^* (*fru^△B/-^*) function does not. (B-D) Overexpressing 2 copies of a *UAS-mam^DN^* transgene in immature INPs partially suppresses the supernumerary neuroblast phenotype in *numb^hypo^* brains. (E) Loss of *fru^C^* function (*fru^△C/-^*) enhances the supernumerary neuroblast phenotype in *brat^hypo^* brains heterozygous for a *fru* deletion (*fru^-/+^)*, but loss of *fru^A^ (fru^△A/-^*) or *fru^B^* (*fru^△B/-^*) function does not. Scale bars: 10 μm. P-value: NS: non-significant, *<0.05, **<0.005, ***<0.0005, and ****<0.00005.

Because Fru^C^ occupies enhancers relevant to *Notch* expression in type II neuroblasts (Fig. 3C), we assessed whether Fru^C^ fine-tunes Notch expression during asymmetric neuroblast division. Proteolytic cleavage of the extracellular domain and the transmembrane fragment releases the Notch intracellular domain to form a transcriptional activator complex by binding Su(H) and Mastermind (Mam) (Bray and Gomez-Lamarca, 2018). Asymmetric segregation of Numb into immature INPs inhibits continual Notch activation and terminates Notch-activated transcription of its downstream-effector genes. *numb*-hypomorphic (*numb^hypo^*) animals carrying the *numb^Ex^* allele *in trans* with a *numb*-null allele (*numb^15^*) contain more than 100 type II neuroblasts per brain lobe compared with 8 per lobe in wild-type animals (Fig. S3A–B). Antagonizing Notch-activated gene transcription by overexpressing a dominant negative form of Mam (Mam^DN^) in immature INPs suppressed the supernumerary neuroblast phenotype in *numb^hypo^* brains (Fig. 4B–D, S3B). Thus, the supernumerary type II neuroblast phenotype in *numb^hypo^* brains provides a direct functional readout of activated Notch levels during asymmetric neuroblast division. The heterozygosity of a *fru* deletion alone did not affect INP commitment in immature INPs but led to a two-fold increase in supernumerary neuroblasts in *numb^hypo^* brains (Fig. 4E). Complete loss of *fru^C^* function (*fru^-/△C^*) led to increased supernumerary neuroblast formation in *numb^hypo^* brains compared with *fru^-/+^* (Fig. 4E). These results suggest that loss of *fru^C^* function increases Notch levels in mitotic neuroblasts leading to higher levels of activated Notch in immature INPs. Thus, we conclude that Fru^C^ fine-tunes *Notch* and Notch downstream-effector gene expression during asymmetric neuroblast division.

### Fru^C^ fine-tunes gene expression by promoting low-level H3K27me3 enrichment

Insights into Fru^C^-mediated fine-tuning of Notch pathway gene expression will likely provide a generalizable mechanism by which Fru^C^ regulates the transcription of thousands of genes during larval brain neurogenesis. A previous study suggested that Fru^C^ functions through Histone deacetylase 1 (Hdac1) to regulate gene transcription during specification of sexually dimorphic neurons (Ito et al., 2012). Acetylated lysine 27 on histone H3 (H3K27ac) is one of the best-characterized residues regulated by Hdac1. We hypothesized that if Fru^C^ fine-tunes *Notch, dpn* and *E(spl)mγ* expression in type II neuroblasts by promoting deacetylation of H3K27 at their promoters and neuroblast-specific enhancers, these loci should display higher H3K27ac levels in *fru^C^*-null brains than in control brains. To ensure the homogeneity of cell types, we compared genome-wide patterns of histone marks by CUT&RUN in *brat*-null brains carrying a *fru^-/△C^* allelic combination (*fru^C^*-null) with that of *brat*-null brains alone (control) (Fig. S4A–B). Fru^C^-bound *cis*-regulatory elements and the bodies of *Notch, dpn* and *E(spl)mγ* display similar patterns of H3K27ac in control and *fru^C^*-null brains (Fig. 5A). To unbiasedly assess whether Fru^C^-bound peaks show increased H3K27ac levels, we identified all 500-bp regions in the entire fly genome that show greater than 2-fold changes in H3K27ac levels in *fru^C^*-null brains relative to control brains (Fig. S4C). A small fraction of regions that show greater than a 2-fold increase in H3K27ac levels overlaps with Fru^C^-bound peaks, but these regions do not include *Notch* or Notch downstream-effector genes that promote stemness in neuroblasts (Fig. S4D). Thus, histone deacetylation likely plays a minor role in Fru^C^-mediated negative regulation of target gene expression. H3K4me3 is a chromatin mark associated with the promoters of actively transcribed genes (Cenik and Shilatifard, 2021). We applied a CUT&RUN assay in control and *fru^C^*-null brains to determine whether Fru^C^ negatively regulates gene transcription by promoting demethylation of H3K4me3. We found that Fru^C^-bound promoters and the bodies of *Notch, dpn*, and *E(spl)mγ* display similar H3K4me3 patterns in control and *fru^C^*-null brains (Fig. 5A). Although many regions that show greater than 2-fold changes in H3K4me3 levels in *fru^C^*-null brains relative to control brains overlap with Fru^C^-bound peaks, their correlation is inconsistent with Fru^C^ negatively regulating gene transcription in type II neuroblasts (Fig. S4E,F). Thus, it is unlikely that Fru^C^ negatively regulates gene expression by promoting demethylation of H3K4me3. These data strongly suggest that Fru^C^ does not finely tune gene expression by regulating the enrichment of active histone marks.

**Fig. 5.**
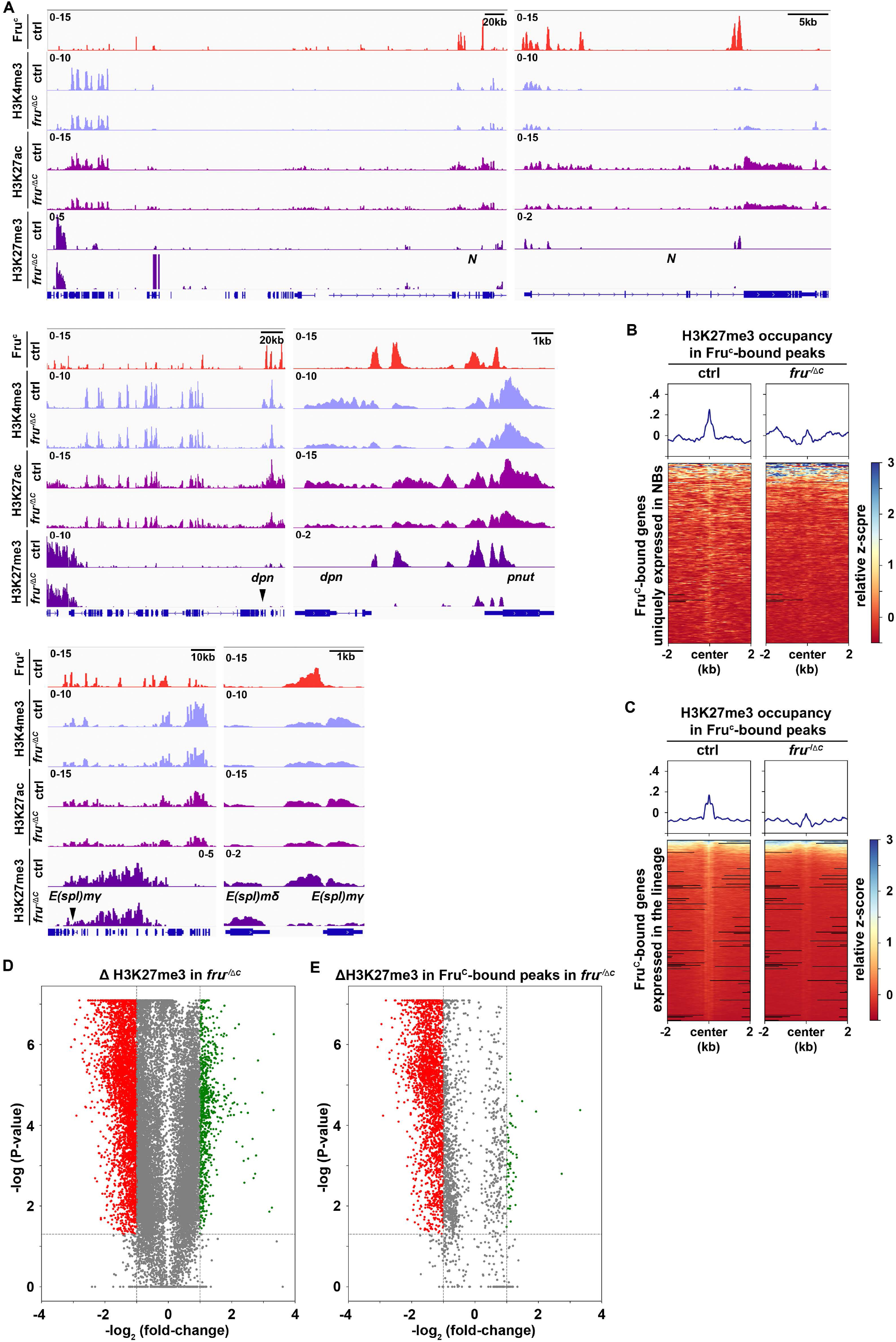
Low levels of H3K27me3 are enriched in Fru^C^-bound regions. (A) Representative z score-normalized genome browser tracks showing Fru^C^-binding and the enrichment of H3K4me3, H3K27ac, and H3K27me3 at *Notch, dpn*, and *E(spl)mγ* loci in type II neuroblasts in control (*brat^-/-^*) or *fru^-/-^* (*brat^-/-^*; *fru^△C/-^*) brains. Left: Zoomed-out images showing nearest heterochromatin domains. Right: Zoomed-in images showing enrichment of histone marks in Fru^C^-bound regions. (B) Heatmaps is centered on Fru^c^ summits with 2-kb flanking regions in genes uniquely transcribed in type II neuroblasts in control or *fru^C^*-null brains and ordered by signal intensity of H3K27me3 enrichment calculated from TMM-normalized tracks. (C) Heatmaps is centered on Fru^c^ summits with 2-kb flanking regions in genes transcribed throughout the type II neuroblast lineage in control or *fru^C^*-null brains and ordered by signal intensity of H3K27me3 enrichment calculated from TMM-normalized tracks. (D) Volcano plot showing fold-change of H3K27me3 signal in 500bp regions in *fru^△C/-^* brains vs control. (E) Volcano plot showing fold-change of H3K27me3 signal in 500bp regions bound by Fru^c^ in *fru^△C/-^* brains vs control.

High levels of H3K27me3 are associated with inactive enhancers and the body of repressed genes (Laugesen et al., 2019; Piunti and Shilatifard, 2021). Low H3K27me3 levels occur in active loci in human and mouse embryonic stem cells, but are generally regarded as noise (Mikkelsen et al., 2007; Pan et al., 2007). If Fru^C^ negatively regulates *Notch, dpn*, and *E(spl)mγ* transcription in type II neuroblasts by promoting H3K27me3 enrichment at their promoters and enhancers, these loci should display lower H3K27me3 levels in *fru^C^*-null brains compared with control brains. While low levels of H3K27me3 is detected in Fru^C^-bound peaks in *Notch* and Notch downstream-effector gene loci in control brains, H3K27me3 levels at these peaks decline significantly in *fru^C^*-null brains (Fig. 5A). We took an unbiased approach to examine whether loss of *fru^C^* function reduces H3K27me3 levels at all Fru^C^-bound genes uniquely transcribed in type II neuroblasts (Fig. 3D). We found that Fru^C^-bound peaks in these genes show lower levels of H3K27me3 enrichment in *fru^C^*-null brains than in control brains (Fig. 5B). Similarly, loss of *fru^C^* function reduces H3K27me3 enrichment at all Fru^C^-bound genes transcribed throughout the type II neuroblast lineage (invariant genes) (Fig. 5C, 3D). Furthermore, many genomic regions that show greater than a 2-fold decrease in H3K27me3 levels in *fru^C^*-null brains relative to control brains overlap with Fru^C^-bound peaks (Fig. 5D–E). These data suggest that Fru^C^ promotes low levels of H3K27me3 enrichment in Fru^C^-bound peaks in genes transcribed in type II neuroblasts. Thus, we conclude that Fru^C^ fine-tunes stemness gene expression by promoting low-level repressive histone marks at their *cis*-regulatory elements in neuroblasts.

### PRC2 fine-tunes gene expression during asymmetric neuroblast division

PRC2 is thought to be the only enzymatic complex that catalyzes H3K27me3 deposition (Laugesen *et al*., 2019; Piunti and Shilatifard, 2021). If Fru^C^ functions through low levels of H3K27me3 enrichment to finely tune *Notch, dpn* and *E(spl)mγ* expression in mitotic neuroblasts, PRC2 core components should be enriched in Fru^C^-bound peaks in type II neuroblasts and reducing PRC2 activity should enhance the supernumerary neuroblast phenotype in *numb^hypo^* brains, identical to the result obtained by reducing *fru* function. Suppressor of zeste 12 (Su(z)12) and Chromatin assembly factor 1, p55 subunit (abbreviated as Caf-1) are two of the PRC2 core components. We performed CUT&RUN in control brains to determine whether regions enriched with Su(z)12 and Caf1 overlap with Fru^C^-bound peaks. We found that Fru^C^-bound *cis*-regulatory elements in *Notch, dpn*, and *E(spl)mγ*display enrichment of Su(z)12 and Caf1 (Fig. 6A). Furthermore, most Fru^C^-bound peaks in genes uniquely transcribed in type II neuroblasts show Su(z)12 and Caf1 enrichment (Fig. 6B). Similarly, most Fru^C^-bound peaks in genes transcribed throughout the type II neuroblast lineage also shows Su(z)12 and Caf1 enrichment (Fig. 6C). These data support our model that Fru^C^ functions through low levels of H3K27me3 enrichment to finely tune gene transcription in type II neuroblasts. Reducing PRC2 function alone does not lead to supernumerary neuroblast formation but strongly enhances the supernumerary neuroblast phenotype in *numb^hypo^* brains, identical to the results obtained by reducing *fru* function (Fig. 6D–F, S4A). Thus, reducing PRC2 activity increases activated Notch during asymmetric neuroblast division. These data led us to propose that Fru^C^ functions together with PRC2 to finely tune gene expression in mitotic neuroblasts by promoting low-level enrichment of repressive histone marks in *cis*-regulatory elements.

**Fig. 6.**
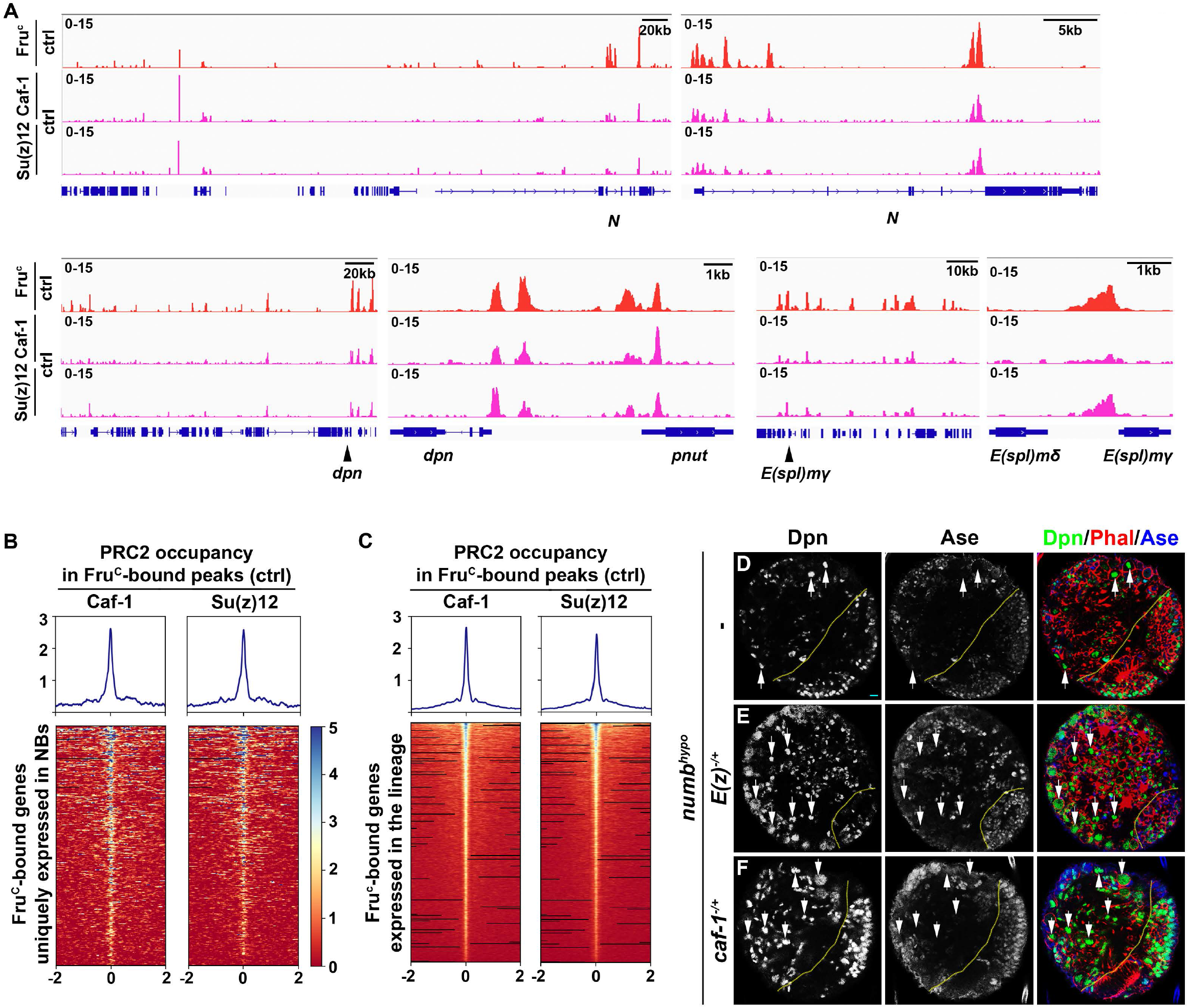
PRC2 subunits bind a high percentage of neuroblast-specific genes. (A) Representative z score-normalized genome browser tracks showing regions bound by Fru^C^, Su(z)12, and Caf-1 in type II neuroblasts in control (*brat^-/-^*) brains. Left: Zoomed-out images of the loci. Right: Zoomed-in images showing enrichment of PRC2 subunits in Fru^c^-bound regions. (B-C) Heatmaps are centered on Fru^c^ summits with 2-kb flanking regions in genes uniquely transcribed in type II neuroblasts in control brains and ordered by average signal intensity of Caf-1 and Su(z)12. Heatmap intensity is calculated from z score-normalized tracks. (D-F) The heterozygosity of E(z) (*E(z)^-/+^*) or caf-1 (*caf-1^-/+^*) enhances the supernumerary neuroblast phenotype in *numb^hypo^* brains. Scale bars: 10 μm.

## Discussion

Regulation of gene expression requires transcription factors and their associated chromatin-modifying activity, and a lack of insights into relevant transcription factors has precluded mechanistic investigations of gene expression by fine-tuning. By taking advantage of the well-established cell type hierarchy and sensitized genetic backgrounds in the type II neuroblast lineage, we demonstrated Fru^C^ fine-tunes gene expression in neuroblasts. By focusing on the Notch signaling pathway in type II neuroblasts, we have been able to define generalizable mechanisms by which Fru^C^ finely tunes gene expression levels using loss- and gain-of-function analyses. Our data indicate that Fru^C^ likely functions together with PRC2 to dampen the expression of specific genes in mitotic neuroblasts by promoting low levels of H3K27me3 enrichment at their enhancers and promoters (Fig. 7). We propose that local low-level enrichment of repressive histone marks can act to fine-tune gene expression.

**Fig. 7.**
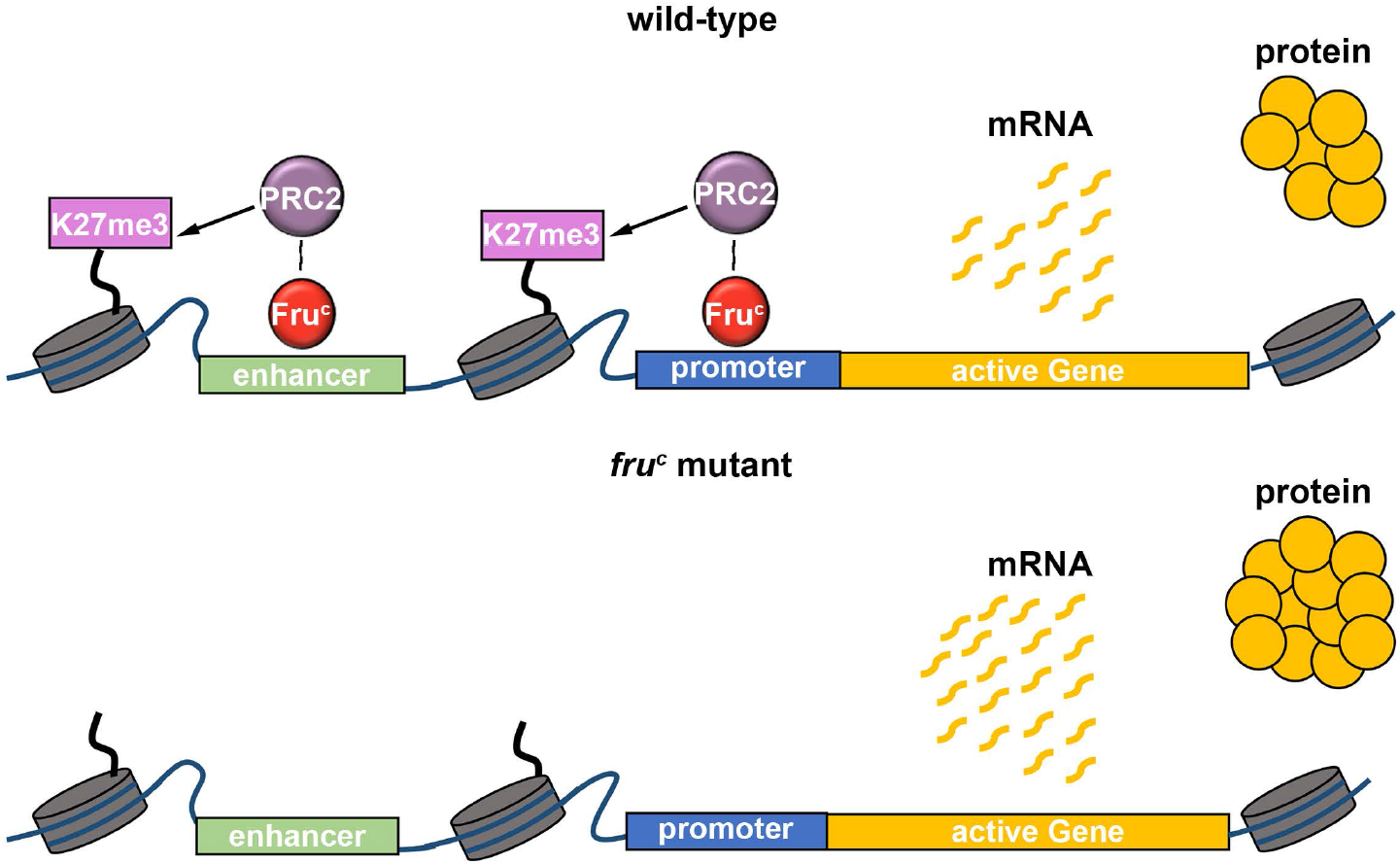
Model. Fru^C^ likely functions together with PRC2 to dampen the expression of stemness genes by promoting low levels of H3K27me3 at their *cis*-regulatory elements. Loss of *fru^C^* functions leads to reduced repressive histone marks and increased stemness gene expression in neuroblasts.

### Fru is a multifaceted transcriptional regulator of gene expression

Fru protein isoforms have been detected in many cell types in flies (Dillon *et al*., 2022; Djiane et al., 2013; Ito et al., 1996; Michki *et al*., 2021; Ryner et al., 1996; Xu et al., 2022; Zhou et al., 2021). The role of Fru in stem cell differentiation remained undefined. Studies linking Fru isoform-specific DNA-binding across the genome with Fru function have been confounded by multiple issues. These include the co-expression of multiple isoforms with different binding specificities within the same cell types, as well as the heterogeneity and scarcity of cell types which express Fru within the central brain, where male-specific Fru isoforms (Fru^M^) have been most intensely studied (Goodwin and Hobert, 2021). In this study, the identification of Fru^C^ as the sole isoform expressed in type II neuroblasts along with the use of *brat*-null brains, which are highly enrich for these neuroblasts, has enabled the study of Fru^C^ genomic binding in a defined cell type where it is known to be expressed. By using gene activity in the Notch pathway as a *in vivo* functional readout, we have now been able to link genomic and genetic evidence to identify a clear role for Fru^C^ in negatively regulating the expression of Fru^C^-target genes during asymmetric neuroblast division (Fig. 3, 4). Our data strongly correlate the downregulation of *Notch* and Notch downstream-effector gene activity by Fru^C^ to PRC2-mediated low-level enrichment of H3K27me3 (Fig. 5, 6). This mechanistic correlation appears to be broadly applicable to genes that promote stemness or prime differentiation in neuroblasts (Fig. 3D,E). These data led us to propose a model in which Fru^C^ functions together with PRC2 to fine-tune the expression of genes by promoting low-level enrichment of H3K27me3 at their *cis*-regulatory elements. Transcriptomic analyses have revealed that *fru* transcripts are enriched in fly renal stem cells in which Notch signaling plays an important role in regulating their stemness (Xu *et al*., 2022). We speculate that the mechanisms we have described in this study might be applicable to the regulation of gene expression in the renal stem cell lineage. It will also be interesting to investigate whether the male-specific Fru^M^C isoform, having the same DNA-binding specificity, utilizes a similar mechanism to regulate the multitude of developmental programs throughout the brain that contribute to a sexually dimorphic nervous system.

### Mechanisms that fine-tune gene transcription

What mechanisms allow transcriptional factors to promote inactivation of gene transcription vs. the finetuning of their expression? In the type II neuroblast lineage, transcription factors Erm and Hamlet (Ham) function together with histone deacetylases to sequentially inactivate type II neuroblast functionality genes including *tll* and *pointed* during INP commitment (Eroglu et al., 2014; Hakes and Brand, 2020; Janssens *et al*., 2014; Koe et al., 2014; Rives-Quinto *et al*., 2020; Weng et al., 2010; Xie et al., 2016; Zhu et al., 2011). In *erm-* or *ham*-null brains, Notch reactivation ectopically activates type II neuroblast functionality gene expression in INPs, driving their reversion into supernumerary neuroblasts. Importantly, mis-expressing either gene overrides activated Notch activity in neuroblasts, and drives them to prematurely differentiate into neurons. Thus, Erm- and Ham-associated histone deacetylation can robustly counteract activity of the Notch transcriptional activator complex and inactivate Notch downstream-effector gene transcription. In contrast to Erm and Ham, Fru^C^ appears to reduce activity of the Notch transcriptional activator complex instead. Loss of *fru^C^* function modestly increases Notch activity in mitotic type II neuroblasts leading to moderately higher levels of Notch activity and Notch downstream-effector gene expression in immature INPs (Fig. 4A, E). Ectopic Notch activity in immature INPs due to loss of *fru^C^* function can be efficiently buffered by the multilayered gene control mechanism and does not perturb the onset of INP commitment (Fig. 2F–I). Furthermore, neuroblasts continually overexpressing Fru^C^ for 72 hours maintained their identities, despite displaying a reduced cell diameter and expressing markers that are typically diagnostic of Ase^+^ immature INPs (Fig. 2J–L). These results suggest that Fru^C^ dampens rather than overrides activated Notch activity and are consistent with the findings that Fru^C^-bound regions displaying little changes in histone acetylation levels between *fru^C^*-null and control neuroblasts (Fig. S4C–D). Thus, transcriptional repressors that inactivate gene transcription render the activity of transcriptional activators ineffective whereas transcriptional repressors that finetune gene expression dampen their activity.

A key follow-up question on a proposed role for Fru^C^ in fine-tuning gene expression in neuroblasts is the mechanistic link between this transcriptional repressor and the dampening of gene transcription. A previous study suggested that Fru functions through Heterochromatin protein 1a (Hp1a) to promote gene repression during the specification of sexually dimorphic neurons (Ito *et al*., 2012). Hp1a catalyzes deposition of the H3K9me3 mark (Eissenberg and Elgin, 2014). Fru^C^-bound genes in neuroblasts display undetectable levels of H3K9me3, and knocking down *hp1a* function did not enhance the supernumerary neuroblast phenotype in *numb^hypo^* brains (data not presented). Thus, it is unlikely that Fru^C^ finely tunes gene transcription by promoting H3K9me3 enrichment. A small subset of Fru^C^-bound peaks showed increased enrichment of H3K27ac in neuroblasts in *fru^C^*-null brains compared with control brains (Fig. S4D). However, Fru^C^ overexpression mildly reduces activity of the Notch transcriptional activator complex in neuroblasts (Fig. 2J–L). Thus, histone deacetylation appears to play a minor role in Fru^C^-mediated fine-tuning of gene expression in neuroblasts. Most peaks that displayed statistically significantly reduced H3K27me3 levels in *fru^C^*-null neuroblasts compared with control neuroblasts are bound by Fru^C^ (Fig. 5E). Many Fru^C^-bound peaks displayed the enrichment of PRC2 subunits, Su(z)12 and Caf1, and reduced PRC2 function enhanced the supernumerary neuroblast phenotype in *numb^hypo^* brains (Fig. 6D–F, S5A). These results strongly suggest a model in which low-level enrichment of H3K27me3 in *cis*-regulatory elements of Fru^C^-bound genes fine-tunes their expression in neuroblasts (Fig. 7).

### PRC2 fine-tunes gene transcription during developmental transitions

A counterintuitive finding from this study is the role for PRC2 and low levels of H3K27me3 enrichment in fine-tuning active gene transcription in type II neuroblasts. PRC2 is thought to be the only complex that deposits the H3K27me3 repressive histone mark and functions to repress gene transcription (Laugesen *et al*., 2019; Morgan and Shilatifard, 2020; Piunti and Shilatifard, 2021). PRC2 subunits and H3K27me3 are enriched in many active genes in various cell types, including embryonic stem cells and quiescent B cells in mice and human differentiating erythroid cells (Brookes et al., 2012; Frangini et al., 2013; Giner-Laguarda and Vidal, 2020; Kaneko et al., 2013; Morey et al., 2013; Ochiai et al., 2020; Xu et al., 2015). However, the functional significance of their occupancies in active gene loci in vertebrate cells remains unclear due to a lack of sensitized functional readouts and a lack of insight regarding transcription factors for their recruitment. Several similarities exist between these vertebrate cell types and fly type II neuroblasts. First, both vertebrate and fly cells are poised to undergo a cell-state transition. Second, the pattern of PRC2 subunit occupancy in *cis*-regulatory elements of active vertebrate and fly genes appears as discrete peaks that are also enriched with low levels of H3K27me3. Building on the functional evidence collected *in vivo*, we propose that Fru^C^ functions together with PRC2 to fine-tune the expression of genes that promote stemness or prime differentiation by promoting low-level enrichment of H3K27me3 in their *cis*-regulatory elements in type II neuroblasts (Fig. 7). Thus, PRC2-mediated low-level enrichment of H3K27me3 we have described in this study should be broadly applicable to the fine-tuned gene activity attained by dampening transcription during binary cell fate specification and cell-state transitions in vertebrates.

An important question arising from our proposed model relates to how Fru^C^ functions together with PRC2 to dampen gene transcription in neuroblasts. Studies in vertebrates have shown that loss of PRC2 activity mildly increases gene transcription levels, suggesting that PRC2 likely dampens gene transcription (Morey *et al*., 2013; Pherson et al., 2017). A separate study suggested a possible link between PRC2 and transcriptional bursts (Ochiai *et al*., 2020). The transcriptional activator complex binding to promoters affects burst sizes, whereas their binding to enhancers control burst frequencies (Larsson et al., 2019). The dwell time of the transcriptional activator complex bound to *cis*-regulatory elements directly affects the frequency, duration, and amplitude of transcriptional bursts. For example, chromatin immunoprecipitation and live-cell imaging of fly embryos have suggested that the Su(H) occupancy time at *cis*-regulatory elements of Notch downstream-effector genes increases upon Notch activation (Gomez-Lamarca et al., 2018). Our genomic data indicate that Su(H)-bound regions in *Notch* and Notch downstream-effector genes that promote stemness in neuroblasts are bound by Fru^C^ and are enriched for low levels of H3K27me3 and PRC2 subunits (Fig. 3F–G, 5A, 6A). These observations suggest that Fru^C^ and PRC2 might fine-tune expression of Notch pathway components by modulating transcriptional bursts in these loci. Examining levels of Su(H) enrichment at Fru^C^-bound peaks in Notch pathway component loci in *fru^C^*-null neuroblasts relative to control neuroblasts will enable the testing of this model.

## Acknowledgements

We thank Drs. G. Cavalli, J. Kadonaga, J. Lis, and D. Yamamoto for providing us with reagents. We thank the Advance Genomics Core and the Flow Cytometry core for technical assistance. We thank Drs. N. Michki and Y. Li for technical advice and assistance on the single-cell RNA sequencing experiment. We thank Drs. Chris Q. Doe and Derek H. Janssen and members of the Lee lab for helpful discussions, and Science Editors Network for editing the manuscript. We thank the Bloomington *Drosophila* Stock Center, Kyoto Stock Center, and the Vienna *Drosophila* RNAi Center for fly stocks. We thank BestGene Inc. and GenetiVision Corp. for generating the transgenic fly lines. This work is supported by NIH grants R01NS107496 and R01NS111647.

## Material and Methods

### Fly genetics and transgenes

Fly crosses were carried out in 6-oz plastic bottles at 25°C, and eggs were collected in apple caps in 8-hr intervals (4-hr for scRNA-seq). Newly hatched larvae were genotyped and cultured on corn meal caps for 96hrs. For overexpression or knock down experiments, larvae were shifted to 33°C after eclosion to induce transgene expression.

Larvae for MARCM analyses (Lee and Luo, 2001) were genotyped after eclosion and allowed to grow at 25°C for 24hr. Corn meal caps containing larvae were then placed in a 37° C water bath for 90 min to induce clones. Heat-shocked larvae were allowed to recover and grow at 25°C for 72hr prior to dissection.

For CUT&RUN experiments, fly crosses were carried out in 30oz fly condos, and eggs were collected on 10mm apple caps in 12-hr intervals. Newly hatched larvae were genotyped and cultured on corn meal caps for ~5–6 days.

The following transgenic lines were generated in this study: *UAS-fru^C^-myc* and *UAS-ERD::fru^C^^zf^-myc*. The DNA fragments were cloned into *p{UAST}attB*. The transgenic fly lines were generated via φC31 integrase-mediated transgenesis (Bischof and Basler, 2008). *numb^EX112^* alleles were generated by imprecise excision of P{GawB}numb^NP2301^, which was inserted at a P-element juxtaposed to the transcription start site of the *numb* gene. The excised regions were determined by PCR followed by sequencing.

### Immunofluorescent staining and antibodies

Larvae brains were dissected in phosphate buffered saline (PBS) and fixed in 100 mM Pipes (pH 6.9), 1 mM EGTA, 0.3% Triton X-100 and 1 mM MgSO4 containing 4% formaldehyde for 23 min. Fixed brain samples were washed with PBST containing PBS and 0.3% Triton X-100. After removing fixation solution, samples were incubated with primary antibodies for 3 hr at room temperature. After 3 hr, samples were washed with PBST and then incubated with secondary antibodies overnight at 4°C. The next day, samples were washed with PBST and equilibrated in ProLong Gold antifade mount (ThermoFisher Scientific). Antibodies used in this study include chicken anti-GFP (1:2000; Aves Labs, SKU 1020), rabbit anti-Ase (1:400) (Weng *et al*., 2010), rabbit anti-Fru^C^OM (1:500; D. Yamamoto), rabbit anti-Trl (1:500; J.T. Lis), mouse anti-cMyc (1:200; Sigma, SKU: M4439), mouse anti-Su(H) (1:100; Santa Cruz, SKU: 398453), mouse anti-Pros (1:500) (Lee et al., 2006b), rat anti-Dpn (1:1000) (Weng *et al*., 2010), and rat anti-Mira (1:100) (Lee *et al*., 2006b). Secondary antibodies were from Jackson ImmunoResearch Inc and ThermoFisher Scientific. We used rhodamine phalloidin or Alexa Fluor Plus 405 phallloidin (ThermoFisher Scientific) to visualize cortical actin. Confocal images were acquired on a Leica SP5 scanning confocal microscope (Leica Microsystems Inc).

### Quantification and statistical analyses

All biological replicates were independently collected and processed. All statistical analyses were performed using a two-tailed Student’s t-test, and p-values<0.05, <0.005, <0.0005, and <0.00005 are indicated by (*), (**), (***) and (****), respectively in figures. GraphPad Prism was used to generate dot plots.

Dpn was used to identify the type II neuroblast and Mira was used to identify the newly born immature INP nucleus. Pixel intensities of the proteins of interest were measured in the nucleus of the type II neuroblast and corresponding newly born immature INP using Image J software. The relative pixel intensity of the protein of interest in the immature INP was taken in relation to the pixel intensity in the type II neuroblast. All biological replicates were independently collected and processed.

### scRNA-seq of the type II neuroblast lineage

*UAS-dcr2; wor-gal4, ase-gal80; UAS-RFP::stinger* larval brains (n=50) were dissected 96hr after larval hatching in ice cold Rinaldini’s solution during a 45-min interval. Dissected brains were transferred to Eppendorf tubes containing 30μL of Rinaldini’s solution. A total of 10μL of 20mg/mL papain, 10uL of 20mg/mL type-1 collagenase, and 1μL of 15μM ZnCl was added to the tube. Additional Rinaldini’s solution was added to adjust the final volume to be 100 μL. The tube was mixed gently by flicking, and them incubated on a heat block at 37°C for 1-hr, while covered with aluminum foil. During this incubation, the tube was flicked for mixing every 10 min.

After the 1-hr incubation, 5μL of 100μM E-64 was added to stop the papain digestion. Samples were incubated on ice for 2 min, and then centrifuged for 3 min at 500g. Supernatant was carefully removed, and chemical dissociated brains were resuspended in 100μL Schneider’s media with 10% fetal bovine serum (FBS). Mechanical dissociation was performed by setting a P100 pipette to 70μL and titrating 30 times at a frequency of ~1Hz. After titration, cells were diluted with 400μL Schneider’s media with 10% FBS, bringing the total volume to 500μL. A total of 1μL of DRAQ5 DNA stain (ThermoFisher Scientific) was added to label cells apart from debris.

A Sony MA900 FACS machine was used to select for RFP+, DRAQ5+ cells. Cells were sorted into a 1.5mL Eppendorf tube prefilled with 100μL of Schneider’s media with 10% FBS. Approximately 30,000 RFP+, DRAQ5 events were sorted. Cells were transported on ice to the University of Michigan’s Advanced Genomics Core and were loaded for 10X Chromium V3 sequencing following the manufacturer’s instructions.

The mRNA was subsequently reverse-transcribed, amplified, and sequenced on an IlluminaNovaSeq-6000 chip (University of Michigan Advanced Genomics Core). Then, 151-bp paired-end sequencing was performed, with a target of 100,000 reads/cell.

### Data analysis for scRNA-seq

Reads were mapped using Cell Ranger (6.0.1) to the *Drosophila* genome assembly provided by ENSEMBL, build BDGP6.32, with DsRed (Genbank: AY490568) added to the genome.

Downstream scRNA-seq analyses was performed using SCANPY (Wolf et al., 2018). Count matrices were concatenated between our dataset and previously published previously published scRNA-seq data generated from INPs and downstream progeny of the type II lineage (Michki *et al*., 2021). The top 2,000 highly variable genes were identified, and then principal component analysis was performed using these highly variable genes with 50 components. The dataset was then harmonized using Harmony (Korsunsky et al., 2019), and completed in four iterations. Neighborhood identification was computed with k=20, and then UMAP (Becht et al., 2018) was performed using a spectral embedding of the graph. Finally, clusters were identified using the Leiden algorithm (Traag et al., 2019), with a resolution of 1. Cell-type annotation was performed by further clustering of clusters 1 and 14, and labeling was performed based on known marker genes.

Pseudotime analysis was performed by calculating the diffusion pseudotime (Haghverdi et al., 2016) trajectory implementation in SCANPY, using an initial root cell selected as *dpn^+^, pnt^+^, DsRed^+^* and visually based on UMAP (ID: CATTCTAAGCAACTTC).

Differential expression analysis for genes between neuroblasts and immature INPs was completed by separating cluster 14 using Leiden with a resolution of 0.3. SCANPY’s rank_gene_groups was used with method=t-test-overestim_var to determine log-fold changes and adjusted p-values of expressed genes. Differentially expressed genes were defined as having |Fold Change| > 2 and adjusted p-value < 0.05, and invariant genes were defined as having |Fold Change| < 2. Bioinfokit (Renesh Bedre, 2020, March 5) was used to generate the volcano plot figure.

### CUT&RUN on neuroblast-enriched brains

*brat^11/Df^; fru^C^::myc* (control) or *brat^11/Df^; fru^ΔC/^fru^Aj96u3^ (fru^ΔC/-^*) larval brains were dissected in 45-min time windows in PBS and transferred to 0.5mL Eppendorf tubes. Dissected brains were then collected at the bottom of the tube, and supernatant was removed. CUTANA^™^ ChIC/CUT&RUN (Epicypher) was performed per the manufacturer’s protocol, with modifications. Brains were resuspended in 100μL wash buffer, and then homogenized by ~30-50 passes of a Dounce homogenizer. Homogenized samples were transferred to a 1.5mL Eppendorf tube, and then the cells were pelleted by centrifugation (600g for 3min). Next, then pellets were processed using the CUT&RUN kit. A total of 0.5ng of antibody was used per sample, (or 0.5μL if antibody concentration was unknown). A total of 0.5 ng of *Escherichia coli* spikein DNA was added into each sample as a spike-in control.

For control brains, antibodies used were goat anti-cMyc (abcam, ab9132), mouse anti-Su(H), rabbit anti-Trl, rabbit anti-Caf1 (Gift from J. Kadonaga) (Tyler et al., 1996), rabbit anti-Su(z)12 (Gift from G. Cavalli, (Loubière et al., 2016)), rabbit anti-IgG (Epicypher), and rabbit anti-H3K9me3 (abcam, ab8898). For both control and *fru^-/-^* brains, antibodies used were rabbit anti-H3K4me3 (Active Motif: 39159), rabbit anti-H3K27me3 (Sigma Aldrich, 07-449), and rabbit anti-H3K27ac (Active Motif: 39136). A total of 50 brains were collected for transcription factor samples and 25 brains were collected for histone mark samples. All samples were performed in duplicate. Samples targeting acetylation had 100 mM of sodium butyrate (Sigma Aldrich) added to all buffers. Samples using mouse antibodies underwent an additional antibody incubation step, where samples were washed 2x with cell permeabilization buffer after primary antibody incubation, and then were incubated for 1 hr with 0.5ng of rabbit anti-mouse IgG (abcam, ab46540).

Fragmented DNA was diluted to 50μL in 0.1X TE and library prepped using the NEBNext^®^ Ultra^™^ II DNA Library Prep Kit for Illumina^®^ (E7645) with NEBNext^®^ Multiplex Oligos for Illumina (E6440) per the manufacturer’s protocol with modifications. The adaptor was diluted 1:25, and bead clean-up steps were performed using 1.1x AMPure Beads without size selection. The PCR cycle was modified to match specifications provided by the CUTANA^™^ ChIC/CUT&RUN kit. DNA was eluted in 20μL 0.1X TE.

DNA samples were assessed for concentration and quality on an Agilent TapeStation. Samples with greater than 1% adaptor underwent an additional round of bead cleanup. Samples that passed quality control were sequenced on an IlluminaNovaSeq-6000 chip (University of Michigan Advanced Genomics Core). Then, 151-bp paired end sequencing was performed, with a target of at least 10,000,000 reads per replicate.

### CUT&RUN data analysis

Read quality was checked using FastQC (Wingett and Andrews, 2018). Reads were trimmed using cutadapt (Martin, 2011), and aligned to BDGP6.32 using bowtie2 (Langmead and Salzberg, 2012) with the flags –local, --very-sensitive, --no-mixed, --no-discordant –dovetail, and -I 10 -X 700. Samtools (Li et al., 2009) was used to convert file formats and to mark fragments less than 120bp.

For transcription factor samples, only reads with a fragment size <120 bp were kept for downstream analysis. Peaks were called individually on each replicate by MACS version 2 (Zhang et al., 2008), using parameters specified in CUT&RUNTools2 (Yu et al., 2021), and then merged using bedtools (Quinlan and Hall, 2010). Downstream analyses were carried out with both Fru^c^::myc and Fru^COM^ peak sets and showed similar results. The data shown in heatmaps use the Fru^c^::myc peakset. For Fru^c^ peaks, only high confidence peaks (-log_10_(Q-value)>100) were kept for downstream analysis. MACS2 was similarly used to call peaks on H3K9me3 samples, and high confidence peaks (-log_10_(Q-value)>100) were merged using bedtools merge -d 3000. We then blacklisted our transcription factor peak sets against the H3K9me3 peaks to remove heterochromatin regions from the downstream analysis. High signal and low mappability regions defined by the ENCODE Blacklist (Amemiya et al., 2019) were also removed using bedtools. A random peak set was generated by calling bedtools shuffle on the Fru^c^::myc peakset, to generate a background control that covered the same number of regions and same number of bp’s as fruitless.

Peak sets were annotated using HOMER (Heinz et al., 2010), which was used to determine the genomic distribution of the transcription factors and genes associated with each peak. Regulatory regions were defined as peaks in either intronic or intergenic regions. Fru^C^ -bound genes were determined as genes that had a Fru^c^::myc peak annotated as being associated to that gene. These peaks were also classified as immature INP-enriched, invariant, or neuroblast enriched peaks based on their corresponding gene’s classification from our single-cell data. This same process was repeated for the randomized peak set.

Bigwig files for transcription factors were generated using deeptools (Ramírez et al., 2016) bamcoverage with flags –ignoreDuplicates –maxFragmentLength 120 –normalizeUsing RPKM. Correlation between Fru^c^::myc and Fru^COM^ was calculated using deeptools mutiBigWigSummary with default parameters. Correlation was plotted on log-log axes using deeptools plotCorrelation with the flag -log1p and otherwise default parameters. Bigwig tracks were visualized using the Integrative Genomics Viewer (IGV) (Robinson et al., 2011). For data visualization, z score-normalized bigwigs were generated by subtracting the mean read coverage (counts) from the merged replicate read counts in 10 bp bins across the entire genome and dividing by the standard deviation (Larson *et al*., 2021). Heatmaps for Fru^c^::myc signal at open chromatin regions, Su(H) peaks, and Trl peaks were generated using deeptools. Heatmaps for Su(z)12 and Caf-1 were generated at Fru^c^ peaks which are associated with neuroblast enriched genes and for all Fru^c^ peaks.

TMM (Trimmed Mean of M-Values) normalization was performed on histone mark samples to accurately account for differences in library composition and sequencing depth. First, peaks were called on control samples against IgG using GoPeaks (Yashar et al., 2022). featureCounts (Liao et al., 2014) was then used to count reads from each replicate inside the peaks for each histone mark across control and *fru^-/-^* samples. EdgeR (Robinson et al., 2010) calcNormFactors (method = TMM) was called on each histone mark countMatrix, and the final normalization factor was calculated as 1,000,000/(normFactor * number of reads in peaks). Bigwig files were generated for individual replicates by using deeptools bamcoverage on aligned bam files with the –scaleFactor equal to the final normalization factor that was calculated. Bigwigs were then merged using deeptools bigWigCompare, and z score bigwig files for histone marks were generated and visualized in IGV. Heatmaps for H3K27me3 between control (*brat^-/-^*) and (*brat_-/-_*; *fru^ΔC/-^*) were generated at Fru^c^ peaks which are associated with neuroblast enriched genes and for all Fru^c^ peaks.

Differential enrichment for each histone marks between control (*brat^-/-^*) and (*brat^-/-^*; *fru^ΔC/-^*) was performed using diffReps (Shen et al., 2013) with a 500bP sliding window and otherwise default parameters. Bins which overlapped with Fru^c^::myc peaks were marked as Fru^c^ bound. Volcano plots showing -Log_10_(pval-adj) vs Log_2_FoldChange were visualized using bioinfokit (Renesh Bedre, 2020).

### Motif analyses

Motifs were searched for within +/-100b p of Fru^COM^ for *Drosophila* position weight matricies (PWMs) using iCisTarget (Imrichová et al., 2015) with default parameters. Analyses were run using all Fru^COM^ peaks, or only Fru^COM^ peaks associated with promoters or regulatory regions, and the top motif was selected for further analysis. The top motif found using all peaks and using regulatory peaks was the same. The normalized enrichment score (NES, [AUC-μ]/σ was recorded for the motifs. Motif PWMs were obtained and motif locations in the genome were calculated using HOMER scanMotifGenomeWide with a log odd detection threshold of 8. Fru^COM^ and random peaks were extended by +/-200bp and the percent of peaks containing motifs was calculated. *De novo* motif searching was attempted using XSTREME (Grant and Bailey, 2021) and HOMER, but no motifs were confidently identified.

---

**Fig. S1.**
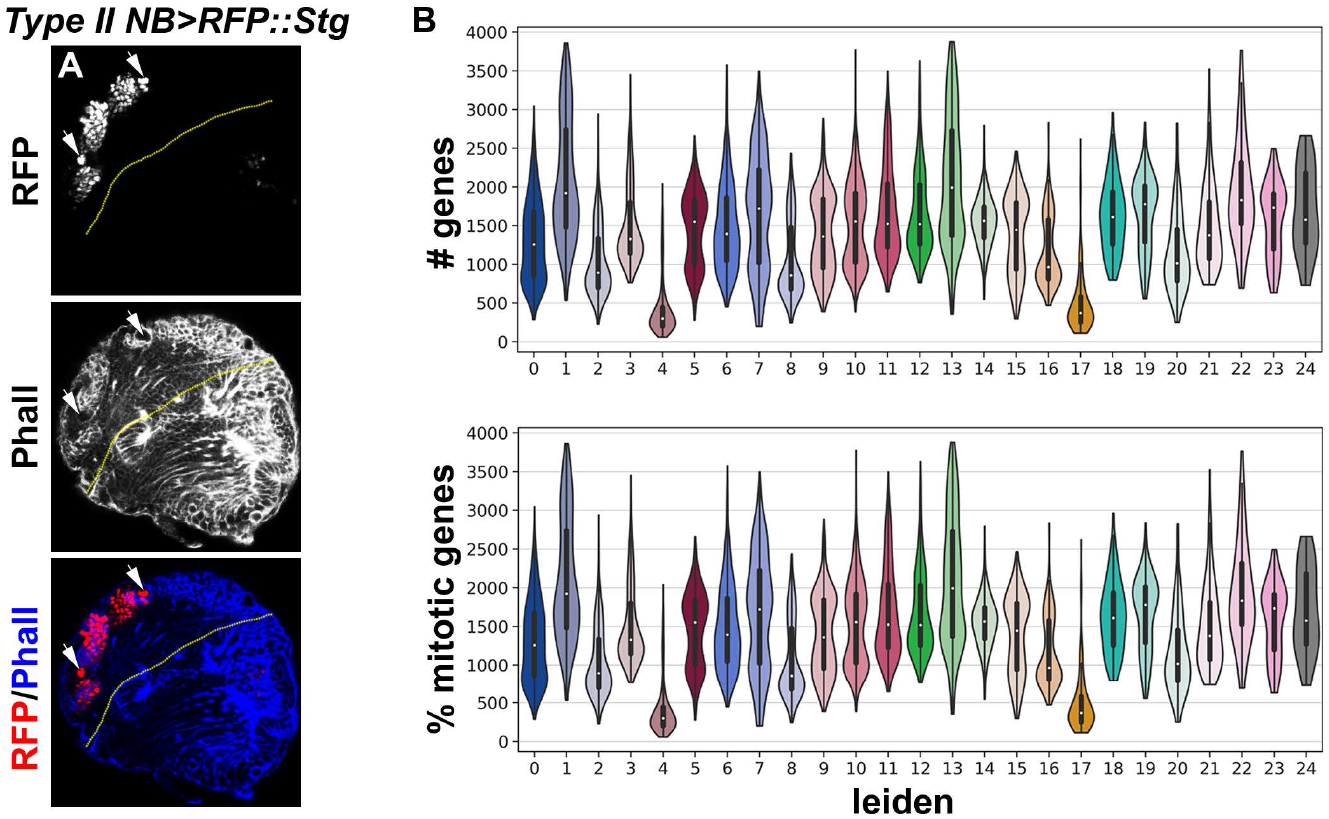
Quality control data for the scRNA-seq atlas. (A) The expression pattern of the *Gal4* driver (*Wor-Gal4,Ase-Gal80*) used in fluorescently labeling and sorting cell types in the type II neuroblast lineage. Endogenously expressed RFP is detectable in the nuclei of all type II neuroblasts and their progeny. (B) Violin plots showing (Top) number of genes or (Bottom) mitochondrial UMI percentage for Leiden clusters shown in 1F. Clusters with low number of genes and higher mitochondrial UMI percentage suggest low-quality or dying cells.

**Fig. S2.**
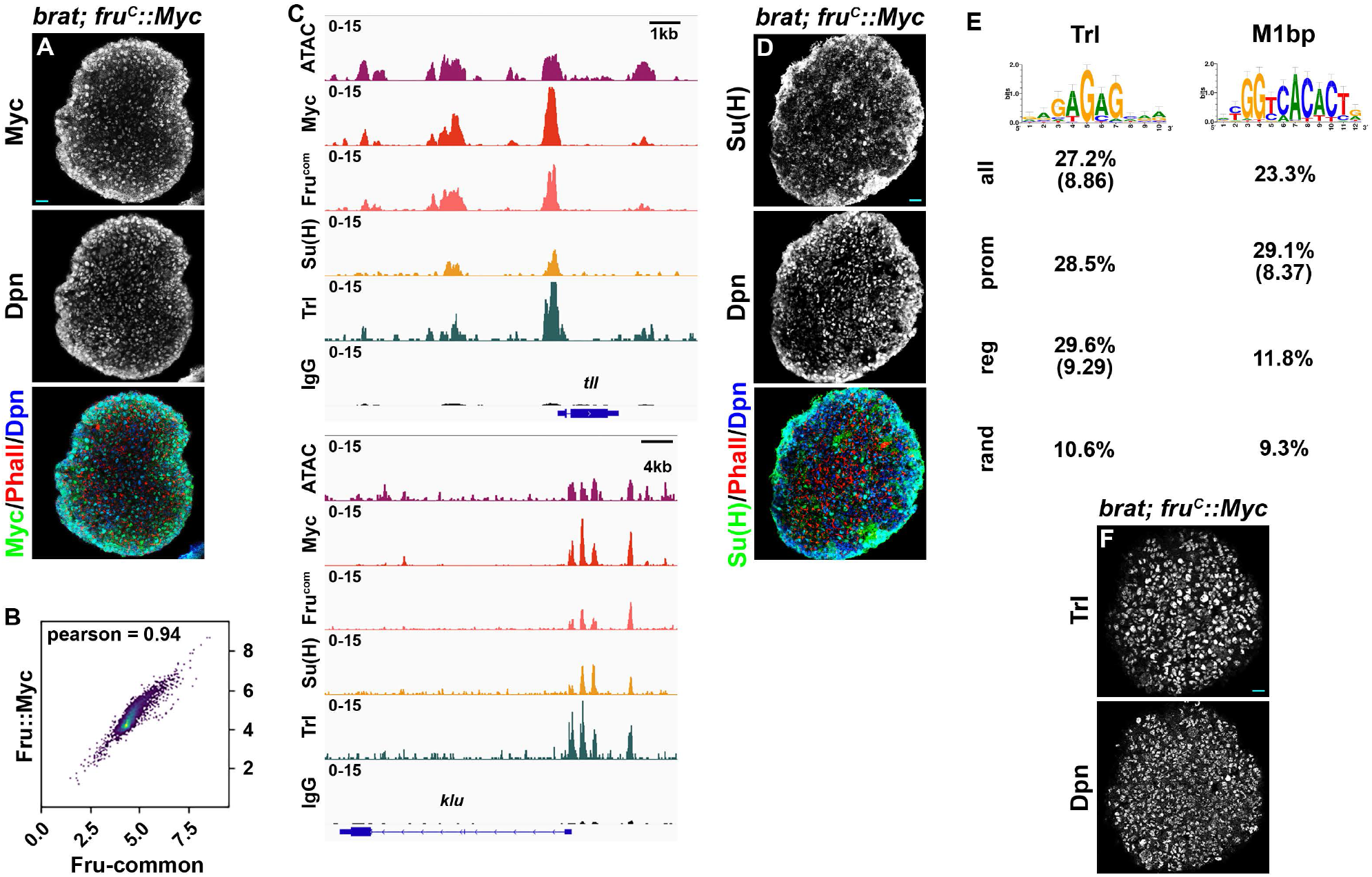
Quality control for determining Fru^c^-, Su(H)- and Trl-binding using CUT&RUN. (A) Fru^C^::Myc is detected in all type II neuroblasts (marked by Dpn expression) in *brat*-null brains homozygous for the *fru^C^::myc* allele. (B) Genome-wide occupancy of Fru^C^::Myc in 10-kb regions in type II neuroblasts determined by the Myc antibody and Fru-common antibody are highly correlated. (C) Representative z score-normalized genome browser tracks showing chromatin accessibility (ATAC-seq) and regions bound by Fru^c^::Myc, Fru^COM^, Su(H), Trl, or IgG at *tll* and *klu* loci. (D) Su(H) is detected in all type II neuroblasts (marked by Dpn expression) in *brat*-null brains homozygous for the *fru^C^::myc* allele. (E) Top motifs identified by iCisTarget from 200-bp regions centered on fru^c^ peak summits. Trl was identified as a top motif using all peaks or only regulator peaks, with NES scores shown in parentheses. M1BP was identified as a top motif using promoter peaks, with NES scores shown in parentheses. The percent of all, promoter, regulatory, or random peaks containing at least one motif was calculated by finding the corresponding motif distribution in the genome with HOMER. (F) Trl is detected in all type II neuroblasts (marked by Dpn expression) in *brat*-null brains homozygous for the *fru^C^::myc* allele. Scale bars: 10 μm.

**Fig. S3.**
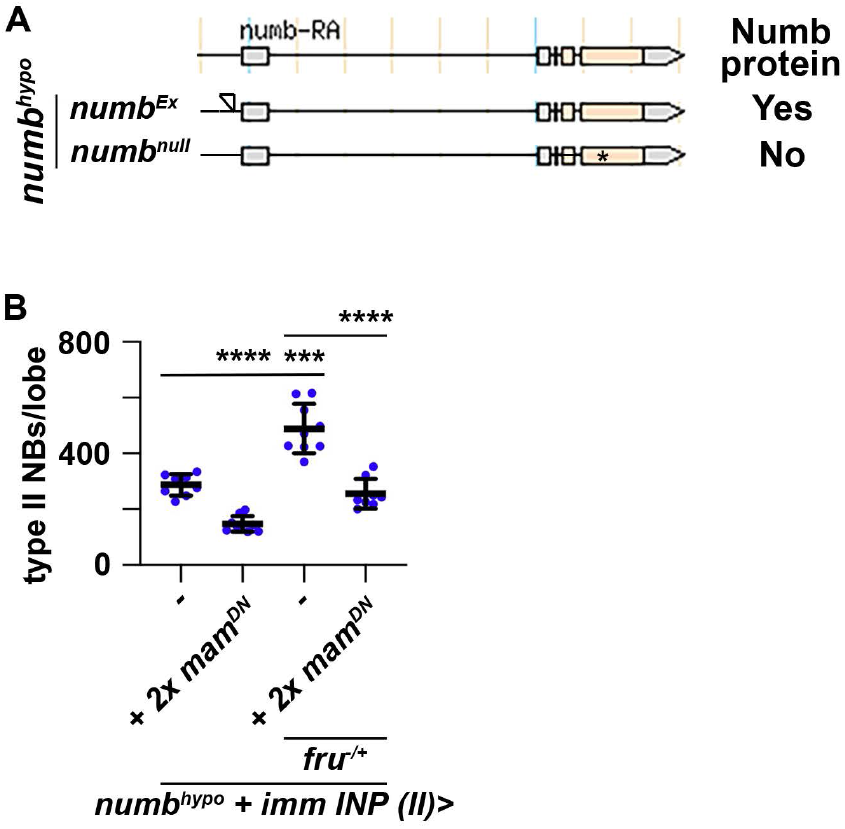
Reduced *fru* function increases Notch downstream-effector gene transcription. (A) The *numb^Ex^* allele was generated by imprecisely excising the *numb^NP2301^* transposable element inserted in the 5’-regulatory region of *numb*. Numb protein remains detectable in *numb^Ex^* homozygous neuroblasts, but is undetectable in *numb*-null neuroblasts. (B) Overexpressing 2 copies of a *UAS-mam^DN^* transgene in immature INPs suppresses the supernumerary neuroblast phenotype in *numb^hypo^* brains induced by the heterozygosity of a *fru* deletion (*fru^-/+^*). P-value: NS: non-significant, *<0.05, **<0.005, ***<0.0005, and ****<0.00005.

**Fig. S4.**
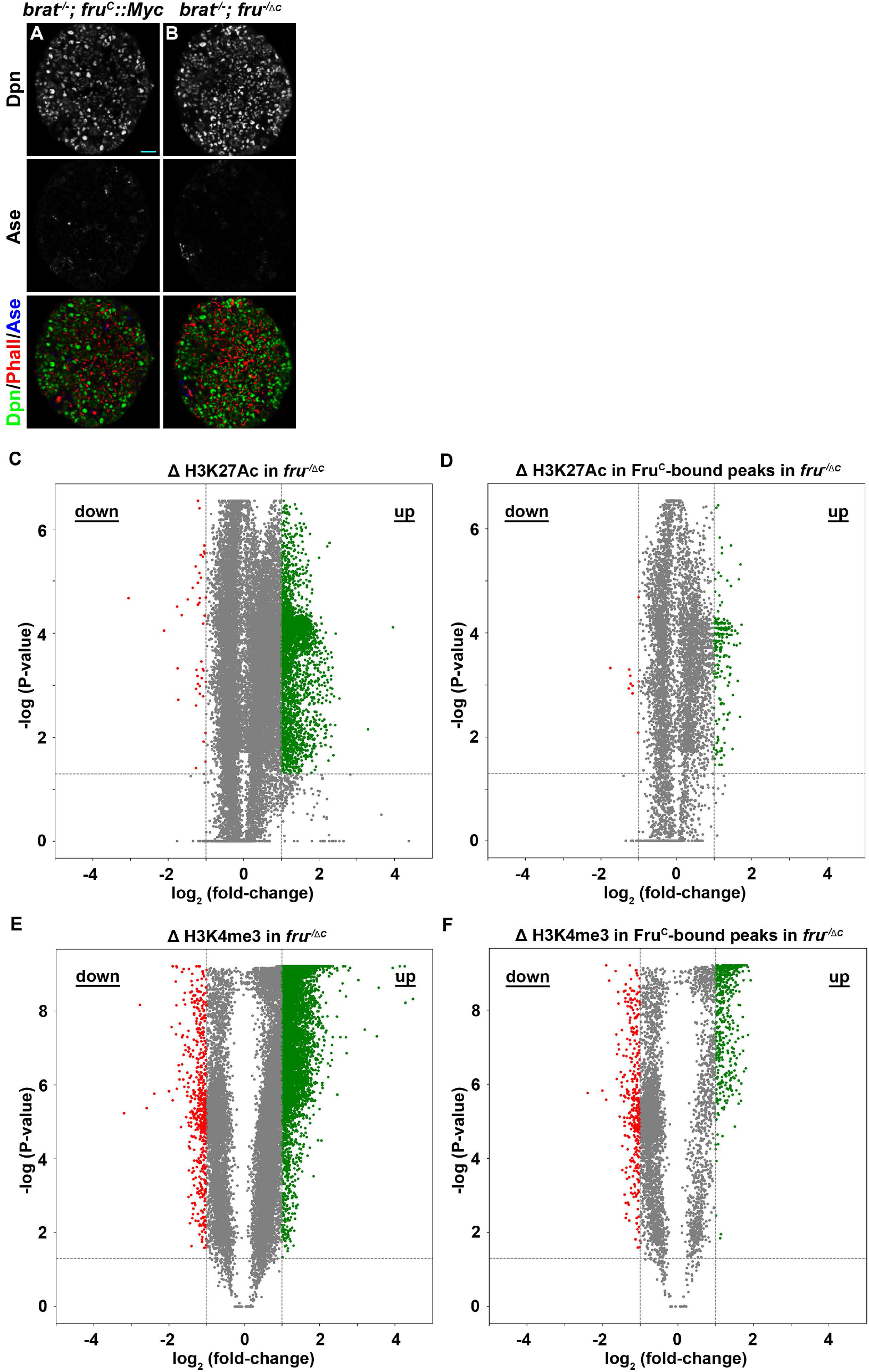
Levels of active histone marks are unchanged Fru^C^-bound regions. (A-B) Loss of *fru^C^* function does not further exacerbate the supernumerary neuroblast phenotype in *brat*-null brains. Scale bars: 10 μm. (C) Volcano plot showing fold-change of H3K27ac signal in 500bp regions in *fru^△C/-^* brains vs control. (D) Volcano plot showing fold-change of H3K27ac signal in 500bp regions bound by Fru^c^. (E) Volcano plot showing fold-change of H3K4me3 signal in 500bp regions in *fru^△C/-^* brains vs control. (F) Volcano plot showing fold-change of H3K4me3 signal in 500bp regions bound by Fru^c^.

**Fig. S5.**
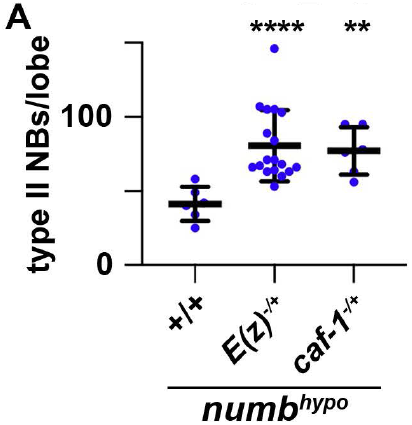
Reduced PRC2 activity enhances stemness gene activity. (A) Quantification of the total type II neuroblasts per brain lobe in *numb^hypo^* brain alone or heterozygous for *E(z)* or *caf-1*.

## Notes

### Competing Interest Statement

The authors have declared no competing interest.

